# Mutations in *VPS18* lead to a neutrophil maturation defect associated with disturbed vesicle homeostasis

**DOI:** 10.1101/2025.08.15.670286

**Authors:** Jincheng Gao, Almke Bader, Linder Monika I., Jingyuan Cheng, Mathis Richter, Annette Zehrer, Karl Mitt, Bastian Popper, Felix Meissner, Megumi Tatematsu, Meino Rohlfs, Stephanie Frenz-Wiessner, Ido Somekh, Joanne Yacobovich, Orna Steinberg-Shemer, Raz Somech, Oliver Soehnlein, Bettina Schmid, Christoph Klein, Barbara Walzog, Daniela Maier-Begandt

## Abstract

Neutrophils, the first cells to arrive at the site of inflammation, are rather short-lived cells and thus have to be constantly replenished. During neutrophil development, vesicle dynamics need to be fine-tuned and impaired vesicle trafficking has been linked to failure in neutrophil maturation. Here, we characterized the role of VPS18 as a central core component of CORVET & HOPS tethering complexes for neutrophil development. Using CRISPR/Cas9-engineered Hoxb8 cells with heterozygous mutations in *Vps18*, we found that VPS18 deficiency interfered with neutrophil development due to tethering complex instability. As a result, vesicle dynamics were impaired with a strong increase in LC3-II and p62 levels, indicating autophagosome accumulation and reduced autophagic flux. With transmission electron microscopy, we verified the increase in autophagosomes and also found irregularly shaped vesicular structures in *Vps18* mutants. Subsequently, *Vps18* mutant neutrophil progenitors underwent premature apoptosis. We described a novel patient with a heterozygous stop-gain mutation in *VPS18* suffering from neutropenia and recurrent infections. To verify our findings in the human system, we used human induced pluripotent stem cells (iPSCs). Upon differentiation into neutrophils, loss of VPS18 resulted in an almost complete absence of iPSC-derived developing neutrophils. Heterozygous *VPS18* mutant and patient mutation-harboring iPSCs were characterized by strongly reduced numbers of developing neutrophils. Zebrafish larvae with heterozygous mutations in *vps18* were also characterized by significantly reduced neutrophil numbers. This study shows the pivotal impact of VPS18 for adequate vesicle dynamics during neutrophil development which might be relevant in the context of vesicle trafficking during granulopoiesis and congenital neutropenia.

## Introduction

Neutrophils are the most abundant leukocytes in human blood and the first cells that arrive at the site of inflammation (1, 2). They are generated during granulopoiesis in the bone marrow with promyelocytes as first committed neutrophil progenitors that give rise to myelocytes, then metamyelocytes, band cells and eventually mature neutrophils. This process is characterized by strong changes in not only nuclear shape, but also production of granules, specific up- and downregulation of cell surface proteins and autophagy-dependent metabolic changes (3–5).

Key for these processes are the fine-tuned intracellular vesicle dynamics which are regulated by an array of proteins. The successful targeting and fusion of vesicles is accomplished by tethering factors that recognize specific proteins on vesicles and target organelles to mediate their spatial contact allowing fusion via soluble N-ethylmaleimide sensitive factor attachment protein receptors (SNAREs) (6). Two tethering factors involved in biosynthetic, endolysosomal and autophagosomal trafficking pathways are class C core vacuole/endosome tethering (CORVET) and homotypic fusion and vacuole protein sorting (HOPS) (7, 8). They consist of six subunits and share the same four core subunits, namely vacuolar protein sorting-associated protein 11 homolog (VPS11), VPS16, VPS18 and VPS33A. CORVET-specific subunits are VPS3 and VPS8 which recognize Ras-related protein Rab-5 (Rab5) to mediate fusion between early endosomes. HOPS specifically harbors VPS39 and VPS41 which bind to Rab7 to allow fusion of lysosomes with late endosomes and autophagosomes, respectively (9).

Patients with homozygous mutations in *VPS16* suffer amongst others from congenital neutropenia together with mucopolysaccharidosis (MPS)-like disease features (10, 11). Patients with homozygous mutations in *VPS33A* were also reported to suffer from congenital neutropenia together with symptoms of an MPS-plus syndrome (12). These data indicate that failure in vesicular trafficking during neutrophil differentiation results in cell death and hence neutropenia. This is in line with findings in congenital neutropenia patients harboring mutations in genes such as *ELANE*, *VPS13B* or *VPS45* where altered vesicle transport leads to apoptosis resulting in neutropenia (13–15).

VPS18 is central for core complex assembly of HOPS by binding to VPS11, VPS16 and VPS41 (16–19). However, the role of VPS18 for neutrophil development is not known to date. *VPS18* knockout (KO) mice die embryonically or early postnatally (20). Mice with a neural-specific *VPS18* KO also die postnatally due to impaired vesicle trafficking and clearance and subsequent neurodegeneration. In zebrafish larvae, mutations of *vps18* result in albinism and liver defects due to impaired endolysosomal trafficking (21, 22).

Here, we show that heterozygous VPS18 deficiency in Hoxb8 cells led to a neutrophil maturation defect followed by premature apoptosis due to CORVET and/or HOPS complex instability and impaired vesicle homeostasis. We further discovered a patient with a novel heterozygous mutation in *VPS18* suffering from neutropenia and recurrent infections. We verified our findings in genetically modified human induced pluripotent stem cells (iPSCs) and *in vivo* in *vps18^+/-^* zebrafish larvae.

## Results

### Deficiency of *Vps18* results in a neutrophil maturation defect

To analyze the role of VPS18 for neutrophil development, we first determined the expression of VPS18 on mRNA and protein level in neutrophils of different origin. We verified the expression of VPS18 in Hoxb8 cell-derived neutrophils (dHoxb8 cells), differentiated HL-60 neutrophil-like (dHL-60) cells and in freshly isolated murine and human neutrophils (Supplementary Fig. 1A). To analyze the role of VPS18 for neutrophil maturation, we employed the murine Hoxb8 cell system. Hoxb8 cells are immortalized neutrophil progenitor cells at the stage of promyelocytes that can be differentiated into mature neutrophils over four days and thus are an excellent model to study neutrophil maturation (23). We generated *Vps18*-deficient Hoxb8 cells using CRISPR/Cas9 technique. Only heterozygous Hoxb8 cell clones were obtained and two different Hoxb8 cell clones were used for our analyses with mutations in the first exon of *Vps18* resulting in a premature stop codon (Supplementary Fig. 1B). The calculated residual protein lengths are 21 (clone 1) and 35 amino acids (clone 2). For control, wild type cells underwent the same procedure but without gRNA and were used as control (CTRL) cells. Downregulation of VPS18 to approx. 50% in both *Vps18* mutants compared to CTRL cells was verified by flow cytometry (Supplementary Fig. 1C-D).

To decipher the potential defect in neutrophil development due to mutations in *Vps18*, Hoxb8 CTRL, clone 1 and 2 cells underwent May-Grünwald-Giemsa staining on each day of differentiation (day 0-4). Promyelocytes, myelocytes, metamyelocytes, band cells and mature neutrophils were quantified based on morphological characteristics (Fig. 1A-B). As described elsewhere, undifferentiated cells at day 0 are mostly in the state of promyelocytes and at day 1 of differentiation develop into myelocytes (24). Initially, CTRL and *Vps18* mutant Hoxb8 cells showed a similar development. From day 2 on, both *Vps18* mutants showed a delayed maturation indicated by reduction of post-mitotic progenitor cells compared to CTRL cells. At day 4 of differentiation, the predominant cell type in CTRL cells were mature neutrophils, whereas in both *Vps18* mutants the majority were dead cells. These data suggest that heterozygous *Vps18* mutations impaired neutrophil development potentially due to insufficient amounts of VPS18. The need of VPS18 for neutrophil development becomes even more evident as VPS18 expression is upregulated during maturation (Supplementary Fig. 1E). Next, essential rescue experiments were performed. Rescue cells were generated by transducing a plasmid coding for the VPS18-EGFP fusion protein into clone 1. Cells were sorted for EGFP fluorescence and analyzed by Western blot for the expression of VPS18-EGFP fusion protein (Supplementary Fig. 1F-G). When these cells underwent differentiation, quantification showed a normal maturation similar to CTRL cells (Fig. 1A-B).

**Figure 1.**
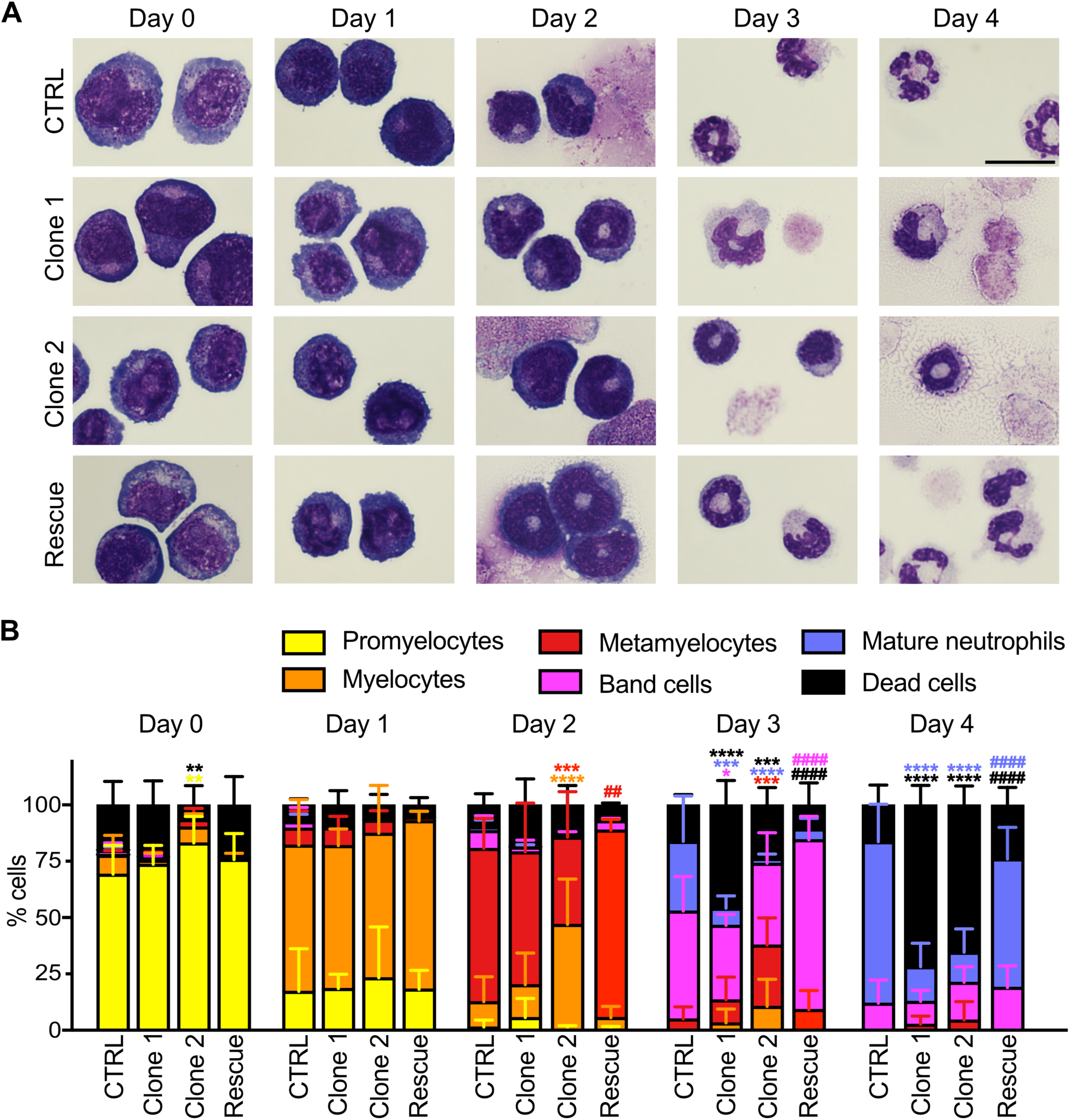
Mutations in *Vps18* cause a maturation defect in neutrophils. (**A**) Representative microscopic images of CTRL, clone 1, clone 2 and VPS18 rescue Hoxb8 cells stained with May-Grünwald-Giemsa during differentiation (day 0-4). Scale bar, 10 µm. (**B**) Quantification of promyelocytes, myelocytes, metamyelocytes, band cells, mature neutrophils and dead cells of indicated genotypes in % (100%, all cells) from (A). n = 5 with a minimum of 200 cells per genotype. Mean ± SD. \**P* < 0.05, \*\**P* < 0.01, \*\*\**P* < 0.001, \*\*\*\**P* < 0.0001 compared to CTRL, ^##^*P* < 0.01, ^####^*P* < 0.0001 compared to clone 1. Two-way ANOVA, Tukey’s multiple comparisons test.

In addition to the morphological-based classification of neutrophil progenitors, a classification system based on cell surface markers has been established (5, 25). When analyzed by spectral flow cytometry and uniform manifold approximation and projection (UMAP), cell clusters during differentiation were visualized and their corresponding maturation stage was identified (Fig. 2A-B). Here, both *Vps18* mutants presented with a late maturation arrest which was characterized by an inability to upregulate the maturation markers CXCR2 and CD101 (Fig. 2C-D). Quantification confirmed the increase in dead cells and the impaired maturation (Fig. 2E; Supplementary Fig. 2). These data verify that neutrophil progenitor cells with mutations in *Vps18* fail to mature normally and instead undergo cell death.

**Figure 2.**
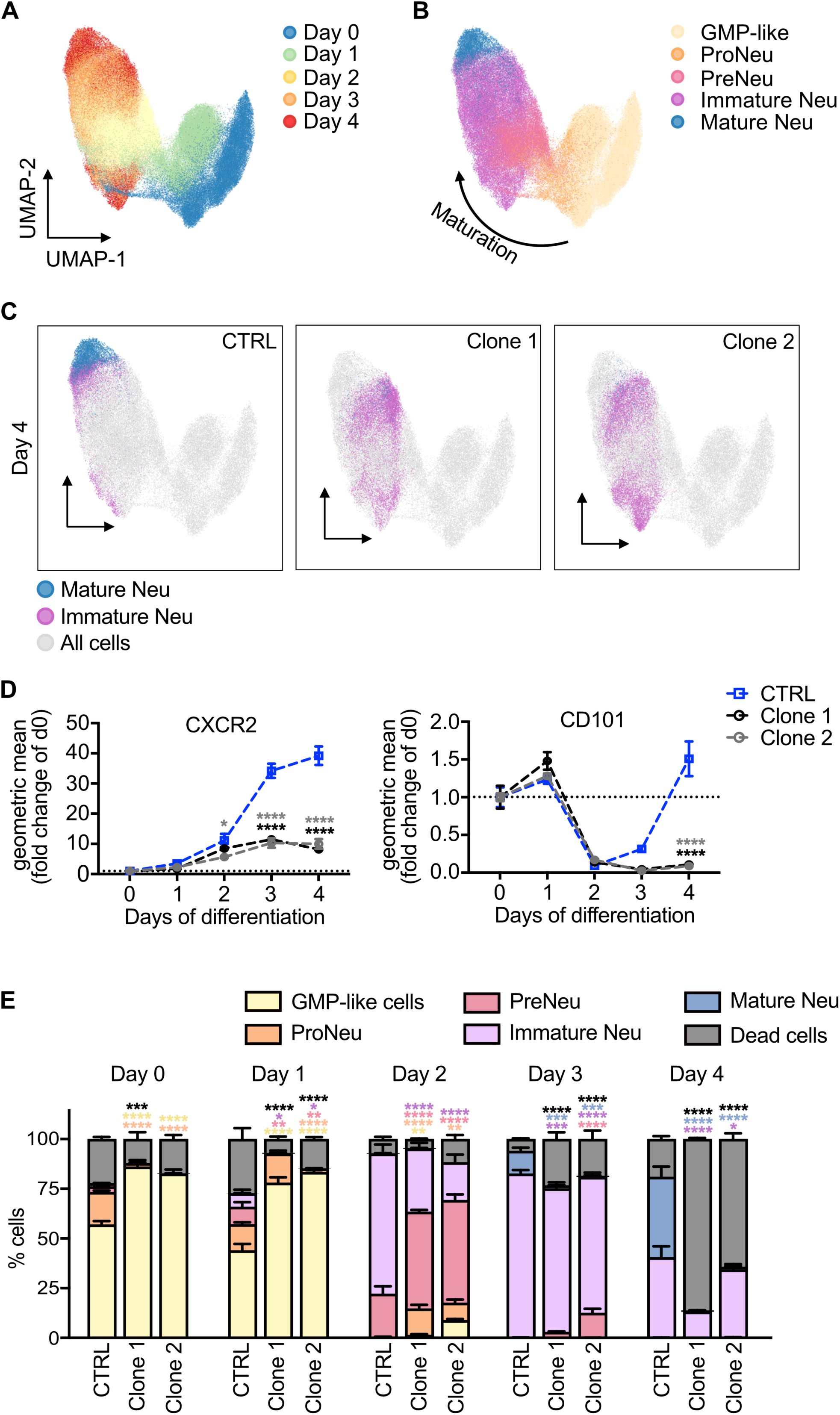
VPS18 deficient neutrophil progenitors have a late maturation defect. (**A and B**) Spectral flow cytometric analysis followed by dimensional reduction using UMAP of all single, living cells at indicated days during differentiation (A), and manually gated differentiation stages (granulocyte-monocyte progenitor (GMP)-like cells, neutrophil progenitors (proNeu), neutrophil precursors (preNeu), immature neutrophils, mature neutrophils) (B). Arrow, direction of maturation. (**C**) UMAP analysis of all single, living cells of CTRL, clone 1 and clone 2 cells at day 4. Clusters of mature (blue) and immature neutrophils (magenta) were manually gated and overlaid onto UMAP plot of all cells during neutrophil differentiation (not gated, grey). (**D**) Relative expression of CXCR2 and CD101 during differentiation (day 0-4) in indicated cell lines using spectral flow cytometry. Geometric mean as fold change of day (d) 0. n = 4. Mean ± SEM. \**P* < 0.05, \*\*\*\**P* < 0.0001 compared to CTRL. Two-way ANOVA, Tukey’s multiple comparisons test. (**E**) Quantification of GMP-like cells, proNeu, preNeu, immature neutrophils, mature neutrophils and dead cells during differentiation from (A-B) of CTRL, clone 1 and clone 2 cells in % (100%, all single cells). n = 4, Mean ± SEM. \**P* < 0.05, \*\**P* < 0.01, \*\*\**P* < 0.001, \*\*\*\**P* < 0.0001 compared to CTRL. Two-way ANOVA, Tukey’s multiple comparisons test.

### *Vps18* deficiency leads to a disturbed intracellular vesicle homeostasis

Next, we wanted to decipher the molecular and cellular effects induced by mutations in *Vps18*. VPS18 is central for tethering complex assembly and we hypothesized that deficiency of *Vps18* led to an instability of the tethering complexes CORVET and HOPS similarly as it was reported for mutations in *VPS16* and *VPS33A* (Fig. 3A) (10, 12). To prove this hypothesis, Hoxb8 cells were analyzed using mass spectrometry (MS) at day 0 and 2 of differentiation (Supplementary Fig. 3A-B). Protein amounts of the direct VPS18 binding partners VPS11 and VPS16 as well as of VPS33A, the other core component, were strongly reduced in *Vps18* mutants compared to CTRL cells at both time points indicating that reduced levels of VPS18 caused an instability of the tethering complexes (Fig. 3B).

**Figure 3.**
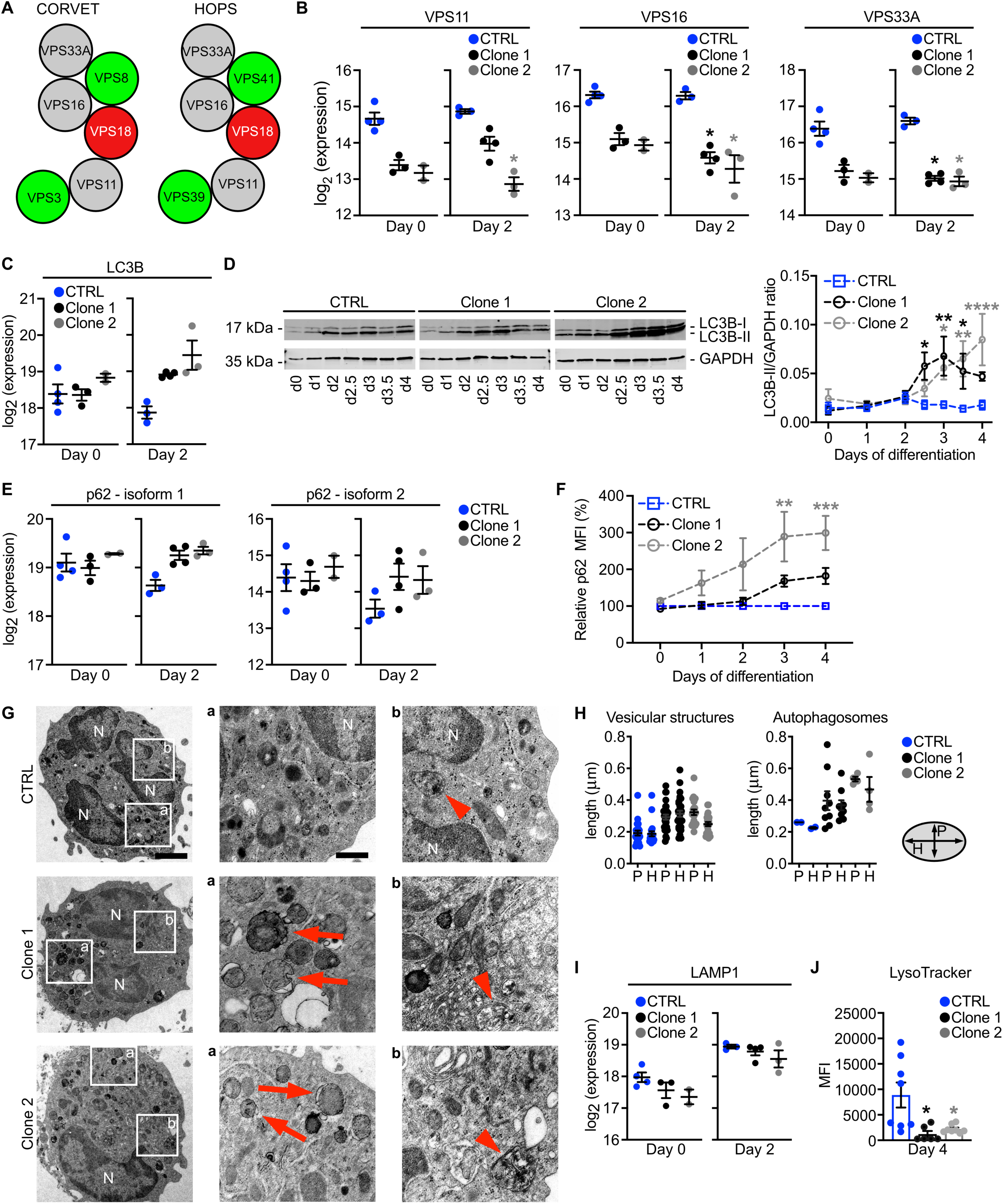
VPS18 deficiency leads to an instability of CORVET and HOPS complexes and to a disturbed vesicle homeostasis during neutrophil maturation. (**A**) Schematic of CORVET and HOPS core (grey) and unique (green) subunits. (**B**) Expression levels of VPS11, VPS16 and VPS33A in indicated cell lines before (day 0) and after removal (day 2) of estrogen analyzed by mass spectrometry. (**C and E**) Expression levels of autophagy markers LC3B and p62 analyzed by mass spectrometry. (**D**) Representative Western blot of LC3B expression (left panel) and quantitative analysis (right panel) of LC3B-II expression in cell lysates from indicated cell lines during differentiation (day 0-4). Ratios of LC3B-II/GAPDH are presented as relative protein amount. n = 7. (**F**) Expression of p62 during differentiation (day 0-4) in indicated cell lines using flow cytometry. Relative p62 levels are shown (normalized to CTRL cells, 100%). n = 9. (**G**) Representative transmission electron microscopy images of CTRL, clone 1 and clone 2 dHoxb8 cells (left panel, scale bar, 2 µm). Squares, zoom-in areas shown in (a) (middle panel, scale bar, 500 nm) and (b) (right panel). Arrows, irregularly shaped vesicular structures. Arrowheads, autophagosomes. N, nucleus. (**H**) Quantification of horizontal (H) and perpendicular (P) size of vesicular structures and autophagosomes from (G). n = 1 independent experiment with n = 5 cells (CTRL & clone 1) and n = 3 cells (clone 2). (**I**) Expression levels of lysosomal marker LAMP1 analyzed by mass spectrometry. (**J**) Quantification of LysoTracker-stained lysosomes in indicated cell lines at day 4. n ≥ 6. (B, C, E, I) Mean ± SEM. n ≥ 2. \**P* < 0.05 compared to CTRL. Pairwise t-test, multiple hypothesis correction. (D and F) Mean ± SEM. \**P* < 0.05, \*\**P* < 0.01, \*\*\**P* < 0.001, \*\*\*\**P* < 0.0001 compared to CTRL. Two-way ANOVA, Tukey’s multiple comparisons test.

Then, the effect of the VPS18 deficiency on the tethering complex target vesicles was investigated. No differences in the amount of RAB5A-C and RAB7, markers for early and late endosomes were found in CTRL cells and *Vps18* mutants (Supplementary Fig. 3C-D). Instead, LC3B and p62, markers for autophagosomes were strongly increased in *Vps18* mutants compared to CTRL cells as detected by MS and verified by Western blot technique indicating an accumulation of autophagosomes and a reduced fusion of autophagosomes with lysosomes (Fig. 3C-F). Using transmission electron microscopy, we found that the size of autophagosomes was increased in *Vps18* mutants compared to CTRL cells accompanied by the appearance of irregularly shaped vesicular structures (Fig. 3G-H). Additionally, the localization of RAB5, RAB7 and LAMP1 as marker for lysosomes shifted to the rear of the cell in adherent *Vps18* mutants compared to their typical localization in the perinuclear center of CTRL cells, further indicating a disturbed vesicle homeostasis in *Vps18* mutants (Supplementary Fig. 3E-F).

We then analyzed levels of LAMP1 and found a trend towards reduced levels in *Vps18* mutants on day 0 and 2 compared to CTRL cells (Fig. 3I). When measuring the fluorescence intensity of acidic compartments in *Vps18* mutant and CTRL cells with LysoTracker, a significant reduction of the signal intensity was present in *Vps18* mutants on day 4 compared to CTRL cells (Fig. 3J). These findings indicate reduced levels of lysosomes which might be due to impaired fusion of lysosomes with target vesicles due to an instability of HOPS.

### *Vps18* mutants display an impaired metabolism and a cell stress response leading to premature apoptosis

Autophagy is induced by various signals to maintain cell homeostasis by for example degradation of damaged intracellular content and supply of metabolic subunits during cell stress such as nutrient starvation (26). In line with this, Riffelmacher et al. have shown that neutrophil development depends on autophagy-mediated lipid breakdown and fatty acid oxidation for mitochondrial respiration and energy production (4). To evaluate the effect of autophagosome accumulation on lipid metabolism in *Vps18* mutants, CTRL and *Vps18* mutant Hoxb8 cells were stained with neutral lipid-specific dyes Nile red and BODIPY (Fig. 4A). Lipid droplet amounts increased in *Vps18* mutant Hoxb8 cells compared to CTRL cells while gene ontology (GO) terms for mitochondrial metabolism showed a decline in *Vps18* mutant Hoxb8 cells as indicated by 1D annotation enrichment analyses (Fig. 4B) suggesting a dysregulated metabolism in *Vps18* mutants. Disturbed vesicle homeostasis and autophagosome accumulation can induce a cell stress response (14, 27). Therefore, we tested for the activation of cell stress markers and found that spliced X-box binding protein 1 (XBP1s) and one of its effector proteins C/EBP homologous protein (CHOP) were significantly increased in *Vps18* mutant Hoxb8 cells compared to CTRL cells (Fig. 4C-D; Supplementary Fig. 4A-B).

**Figure 4.**
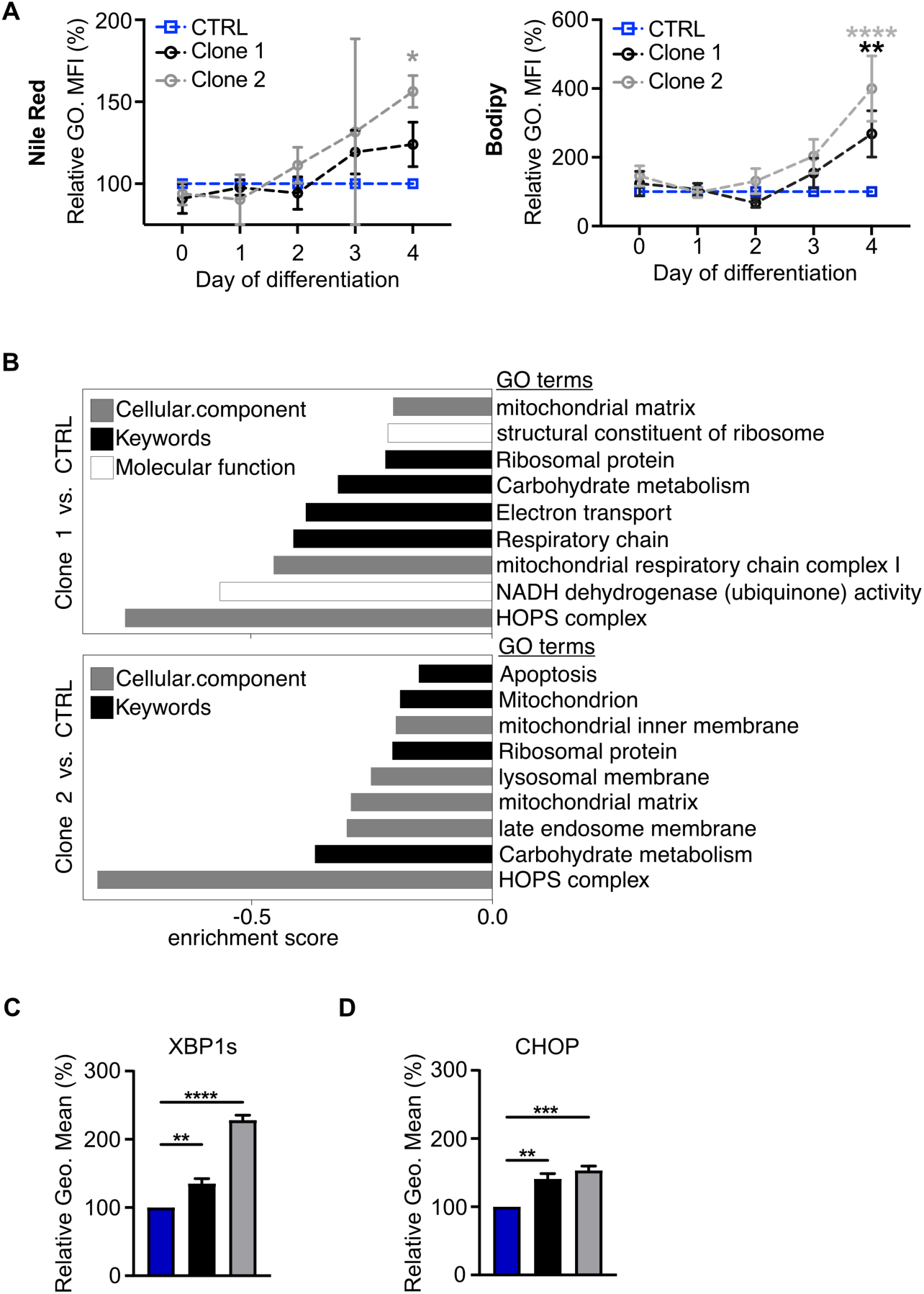
*VPS18^+/-^* mutant Hoxb8 neutrophil progenitors undergo metabolic changes and perceive cell stress. (A) Quantification of lipid-droplet staining with Nile red (upper panel) and Bodipy (lower panel) of indicated genotypes during differentiation (day 0-4) analyzed by flow cytometry (normalized to CTRL cells, 100%). Mean ± SEM. n = 3 (Nile red), n ≥ 7 (Bodipy). \**P* < 0.05, \*\**P* < 0.01, \*\*\**P* < 0.0001 compared to CTRL (d4), two-way ANOVA, Tukey’s multiple comparisons test. (**B**) Strongly reduced gene ontology (GO) terms in clone 1 and 2 compared to CTRL cells at day 2 of differentiation by 1D enrichment analysis of mass spectrometry data. (**C and D**) Quantification of ER-stress markers XBP1s (C) and CHOP (D) at day 3 of differentiation by flow cytometry (normalized to CTRL cells, 100%). n = 4. \*\**P* < 0.01, \*\*\**P* < 0.0001, \*\*\*\**P* < 0.0001 compared to CTRL. Mean ± SEM, one-way ANOVA, Tukey’s multiple comparisons test.

Unresolved cell stress and an impaired autophagosome-lysosome fusion can induce apoptotic cell death (26). Indeed, active caspase-3, an effector caspase during apoptosis was significantly increased in both *Vps18* mutants compared to CTRL cells at day 3 of differentiation (Fig. 5A). This increase was absent in rescue cells compared to clone 1 Hoxb8 cells (Fig. 5B). To further verify the apoptotic cell death that *Vps18* mutants underwent during differentiation, CTRL and *Vps18* mutant Hoxb8 cells were stained with tetramethylrhodamine methyl ester (TMRM), Annexin V and SytoxRed and analyzed by flow cytometry (Fig. 5C-D; Supplementary Fig. 5).

**Figure 5.**
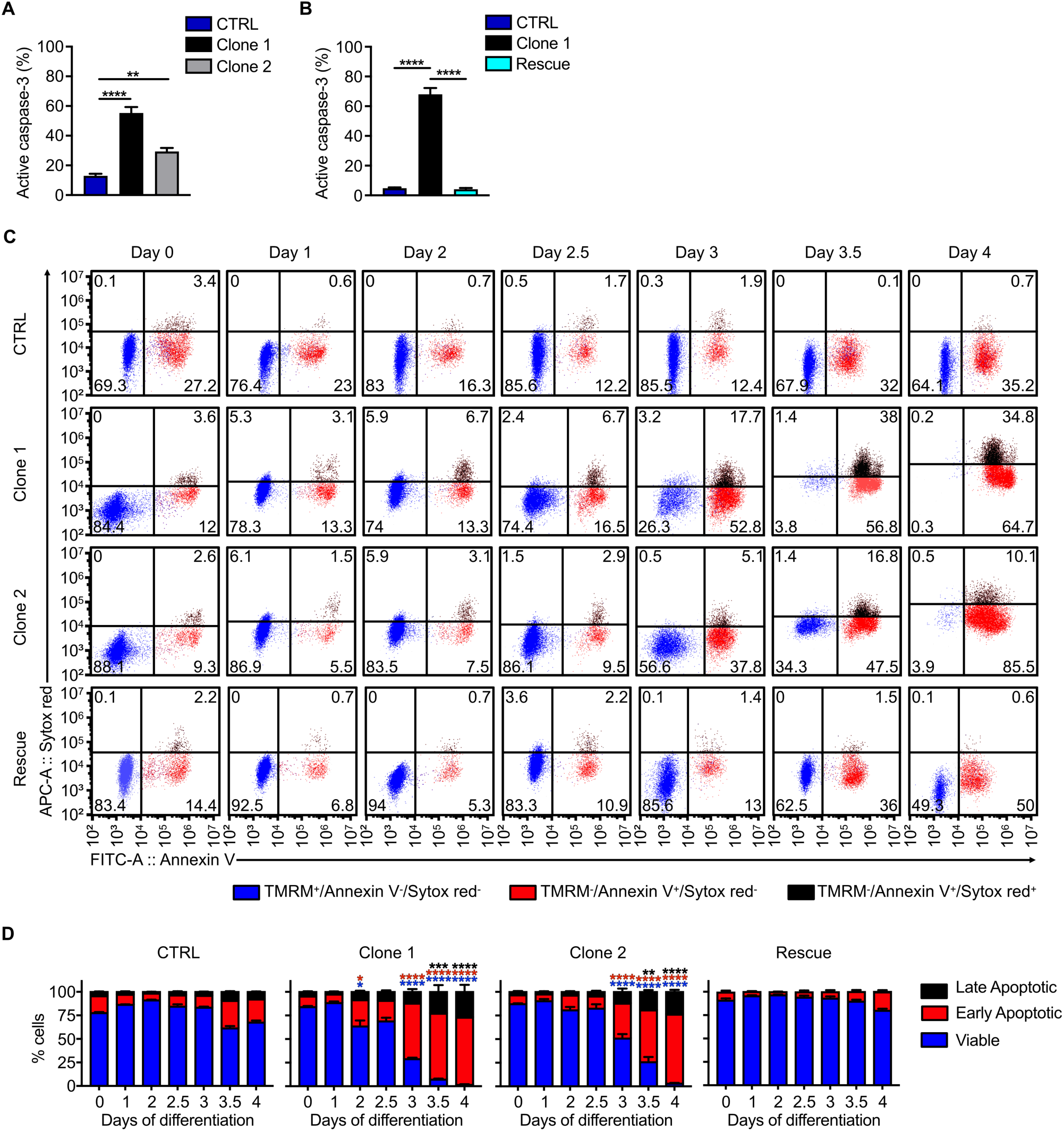
*VPS18^+/-^*mutant Hoxb8 neutrophil progenitors undergo premature apoptosis. (**A and B**) Cells positive for active caspase-3 at day 3 of differentiation in % of all single cells in indicated genotypes analyzed by flow cytometry. n = 6. \*\**P* < 0.01, \*\*\*\**P* < 0.0001 compared to CTRL (A) or compared to clone 1 (B). (**C and D**) Representative dot plots (C) and quantification (D) of viable, early and late apoptotic cells of indicated genotypes during differentiation (day 0-4). Viable cells, TMRM^+^, blue. Early apoptotic cells, Annexin V^+^ and SytoxRed^-^, red. Late apoptotic cells, Annexin V^+^ and SytoxRed^+^, black. Numbers indicate % of cells of all single cells (100%, sum of all single cells). n = 4. \**P* < 0.05, \*\**P* < 0.01, \*\*\**P* < 0.001, \*\*\*\**P* < 0.0001 compared to d0. (A-B, D) Mean ± SEM, one-way ANOVA, Tukey’s multiple comparisons test.

Over four days of differentiation, the majority of CTRL Hoxb8 cells was viable (TMRM^+^) and only 35.2% of CTRL Hoxb8 cells were early apoptotic (Annexin V^+^, Figure 5C-D). In strong contrast, both *Vps18* mutants showed significantly reduced levels of viable cells from day 3-4 accompanied by an increase in early and late apoptotic (Annexin V^+^/SytoxRed^+^) cells. Rescue cells were also analyzed during differentiation and were characterized by the clear absence of apoptotic cell death markers compared to clone 1 Hoxb8 cells. Thus, these data show that heterozygous mutations in *Vps18* induced apoptosis in premature neutrophils which we defined as premature apoptosis.

### Discovery of human VPS18 deficiency

The patient is a 7-year-old female from a non-consanguineous background and was identified during infancy when her symptoms required medical attention. She experienced recurrent respiratory infections of viral and bacterial pathogens including human parainfluenza virus, adenovirus and *Haemophilus influenzae* that required hospitalizations as well as severe diaper rash and recurrent aphthous ulcers. Initial laboratory tests revealed severe neutropenia in serial blood tests during the first year of life, which gradually improved over time (Supplementary Table 1). Anti-neutrophil antibodies were negative, bone marrow aspiration showed an abundance of myelocytes and a reduction in mature neutrophils indicating a late maturation arrest of neutrophils (Fig. 6A). Additional immune workup, including immunoglobulin levels, T- and B-lymphocyte subsets, NK-cell counts, and T-cell receptor excision circles, showed no abnormalities. Genetic investigation employing whole exome sequencing identified a nonsense variant in *VPS18* (c.700C>T; p.Arg234Ter) which was validated by Sanger sequencing (Supplementary Table 2). No revertant mutation was observed. The patient exhibited both a motorical and speech developmental delay during infancy. However, with the resolution of recurrent infections and hospitalizations the patient reached the expected developmental milestones. The patient’s father, which is asymptomatic with normal neutrophil counts was also found to harbor an identical heterozygous variant. The patient is currently in good health with normal neutrophil counts. However, the neutrophil counts of both the patient and the father are in the lower range of the normal counts.

**Figure 6.**
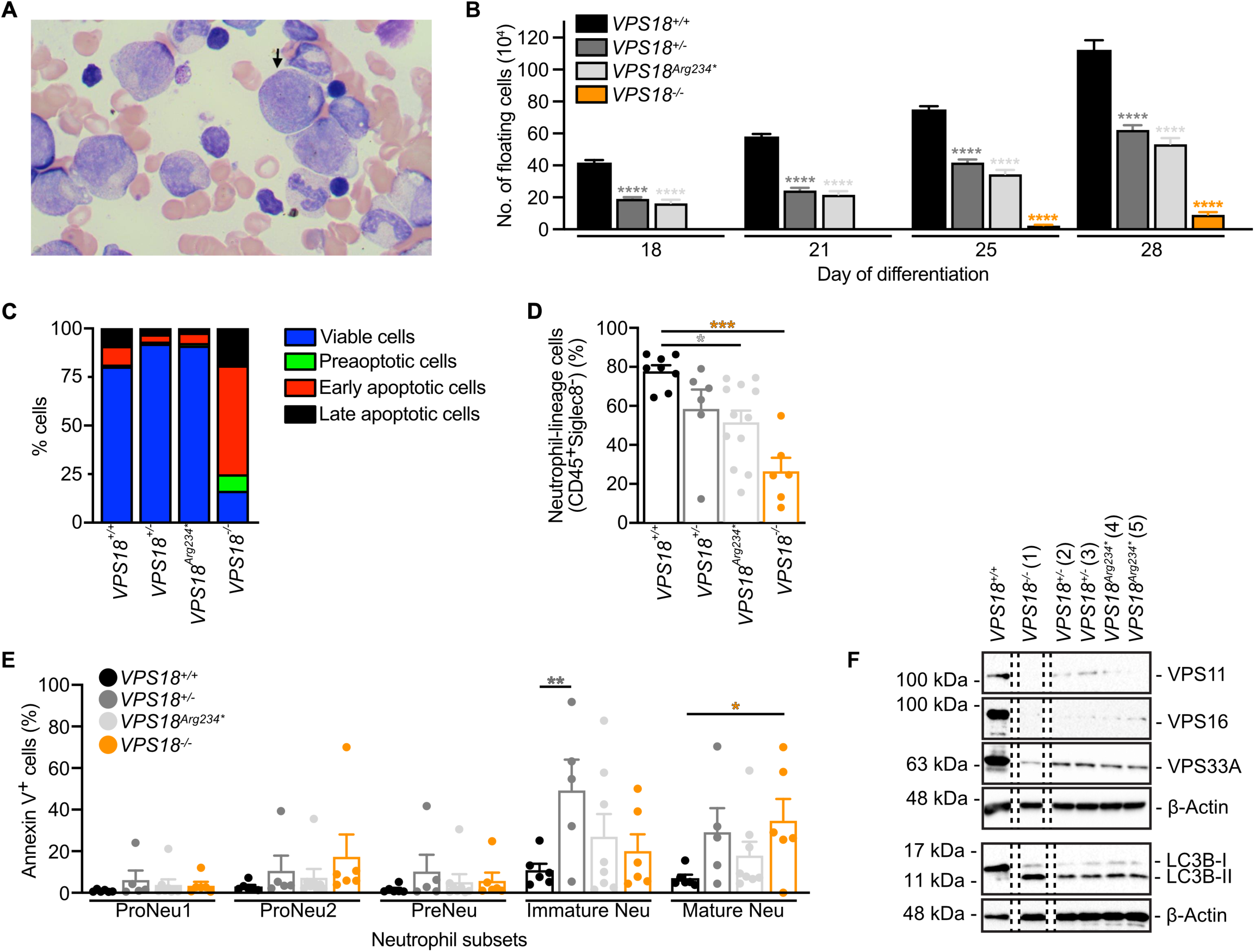
Patient-specific mutations in iPSC-derived neutrophils recapitulate the patient’s phenotype. (**A**) Microscopy image of the patient’s bone marrow aspiration. Myelocyte, arrow. Number of floating cells released from 6 iPSC colonies per well at indicated days of differentiation from indicated cell lines. n = 3 technical replicates per cell lines (1 *VPS18^+/+^*, 2 *VPS18^+/-^*, 2 *VPS18^Arg234*^*, 1 *VPS18^-/-^* cell lines). Mean ± SEM. \*\*\*\**P* < 0.0001 compared to *VPS18^+/+^*. (**C**) Quantification of viable, preapoptotic, early and late apoptotic cells of indicated genotypes at day 28 of differentiation in % (100%, sum of all detected single cells) analyzed by flow cytometry. n ≥ 1. Data presented as mean. (**D**) Quantification of neutrophil-lineage directed cells (CD45^+^Siglec^-^) from indicated cell lines at day 28 analyzed by flow cytometry. N ≥ 6. Mean ± SEM. \**P* < 0.05, \*\*\**P* < 0.001 compared to *VPS18^+/+^*. (**E**) Annexin V^+^ cells within populations of proNeu1 (CD11b^-^, CD49d^+^, SSC-A^low^), proNeu2 (CD11b^-^, CD49d^+^, SSC-A^high^), preNeu (CD49d^+^, CD101^-^), immature (CD35^+^, CD16^-^) and mature (CD35^+^, CD16^+^) neutrophils from indicated cell lines at day 28 in % (100%, sum of all single cells) analyzed by flow cytometry. n ≥ 6. Mean ± SEM. \**P* < 0.05, \*\**P* < 0.01 compared to *VPS18^+/+^*. (**F**) Representative Western blots of VPS11, VPS16 and VPS33A (upper panel) & LC3B expression (lower panel) in cell lysates from indicated cell lines at day 28. β-actin was used as loading control. n = 2. (B-E) Two-way ANOVA, Tukey’s multiple comparisons test.

### Mutations in *VPS18* lead to impaired neutrophil development of human iPSCs and to neutropenia in zebrafish larvae

To verify our data obtained from murine Hoxb8 cells in the human system, we generated human iPSCs with mutations in *VPS18*. Using CRISPR/Cas9 technique, we generated one *VPS18^-/-^* (clone 1), two *VPS18^+/-^* (clone 2 and 3) and two *VPS18^Arg234*^*(harboring the patient-specific mutation, clone 4 and 5) cells (Supplementary Fig. 6A). Loss or a strong downregulation of VPS18 protein was verified by Western blot technique in *VPS18^-/-^* cells or both *VPS18^+/-^*and *VPS18^Arg234*^* cells, respectively (Supplementary Fig. 6B). Addition of hematopoietic cytokines allowed differentiation of iPSCs to neutrophils as described earlier (28). Similar to CTRL Hoxb8 cells, *VPS18^+/+^* iPSCs upregulated VPS18 during differentiation (Supplementary Fig. 6C). Developing neutrophils were released from *VPS18^+/+^*iPSC colonies into the medium resulting in 112.3 x 10^4^ floating cells at day 28 of differentiation (Fig. 6B). In comparison, levels of floating cells were significantly reduced to 50% in *VPS18^+/-^* and *VPS18^Arg234*^* colonies. *VPS18^-/-^* colonies showed an even stronger reduction with 9.3 x 10^4^ floating cells indicating that a full knockout is barely viable. Indeed, bulk analyses at day 28 showed that only 16.3% of *VPS18^-/-^*mutant cells were viable and the majority lost their membrane potential and underwent apoptosis (Fig. 6C; Supplementary Fig. 6D). In comparison, the majority of *VPS18^+/+^*, *VPS18^+/-^* and *VPS18^Arg234*^* cells were viable on this day. We applied an immune phenotype panel as described before and found that the number of cells undergoing neutrophil-lineage differentiation was strongly reduced in *VPS18* mutant cells compared to *VPS18^+/+^* cells (Fig. 6D, Supplementary Fig. 6E-H) (29). The number of cells undergoing apoptosis increased especially in immature and mature neutrophils which is in line with our observations in *Vps18* mutant Hoxb8 cells (Fig. 6E). Levels of VPS18 binding partners VPS11, VPS16 and VPS33A were strongly downregulated in *VPS18* mutant iPSCs compared to *VPS18^+/+^* iPSCs suggesting that VPS18 deficiency and also patient-specific mutations caused HOPS and CORVET complex instability (Fig. 6F). *VPS18* mutant cells were characterized by an increase in LC3B-II compared to *VPS18^+/+^* cells indicating an increase in autophagosomes as seen in *Vps18* mutant Hoxb8 cells.

To verify these data *in vivo*, we employed the zebrafish model. Zebrafish harbor a single orthologue of Vps18 with a sequence identity of 66% and similarity of 79% to human VPS18 (10). Using CRISPR/Cas9 technique, we generated heterozygous *vps18* deficient Tg*(fli1:gfp;lyz:dsRed)* zebrafish targeting exon 1 (*vps18* mutant 1) or exon 4 (*vps18* mutant 2) (Supplementary Fig. 7A). Total neutrophil counts were analyzed at 3 days post fertilization (dpf). Mutations in *vps18* in both mutants resulted in a significant decrease of total neutrophil numbers compared to *vps18* wild-type (WT) zebrafish larvae (Fig. 7). These data are in line with our *in vitro* findings and suggest that already a heterozygous Vps18 deficiency is sufficient for failure of neutrophil development. Interestingly, neutropenia was absent in adult, two-year-old *vps18* mutant zebrafish compared to *vps18* WT zebrafish indicating that mutations in *vps18* might induce a transient neutropenia (Supplementary Fig. 7B-C). We also investigated neutrophil trafficking behavior in a tail fin transection assay in zebrafish larvae and found similar numbers of recruited neutrophils to sites of injury 1, 3, and 6 h post wounding in *vps18* WT and *vps18* mutant 1 zebrafish larvae indicating that Vps18 is dispensable for neutrophil migration (Supplementary Fig. 7D-F).

**Figure 7.**
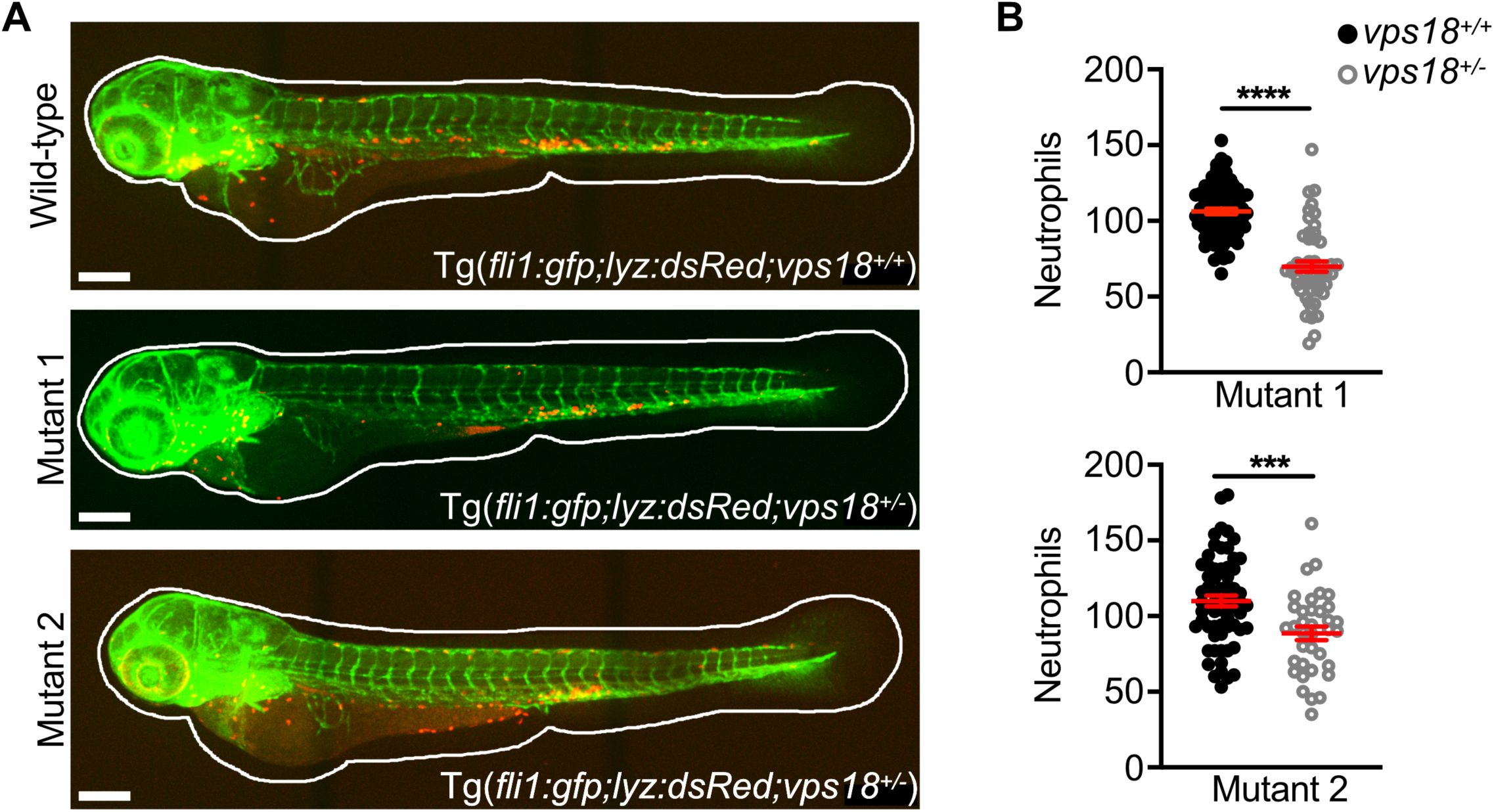
Zebrafish larvae deficient for *vps18* recapitulate the patient’s phenotype. (**A and B**) Exemplary maximum intensity projections (A) and total neutrophil counts (B) of *vps18^+/+^*and *vps18^+/-^* mutant 1 and 2 zebrafish larvae at 3 dpf. Endothelial cells, green. Neutrophils, red. Scale bars, 200 μm. Mean ± SEM of ≥ 36 individual larvae of ≥ 7 independent experiments. \*\*\**P* < 0.001, \*\*\*\**P* < 0.0001. Unpaired t-test.

## Discussion

In this study, Hoxb8 cell, iPSC and zebrafish models were employed to decipher the role of VPS18 for neutrophil development. We found VPS18 to be vital for neutrophil maturation and identified a novel human genetic defect in *VPS18* characterized by neutropenia and a neutrophil maturation defect.

As described, during generation of *Vps18* deficient Hoxb8 cells by CRISPR/Cas9 technique only heterozygous but no homozygous cell clones were obtained, leading to the hypothesis that a complete knockout of *Vps18* at the promyelocyte stage is lethal. Our studies with human iPSCs also indicated that cells with homozygous mutations in VPS18 at the level of pluripotency are viable but are prone to die during differentiation and commitment to the myeloid lineage. In line with these findings Peng *et al.* described that a full *Vps18* KO in mice is embryonically or early postnatally lethal, as is a neural-specific *Vps18* KO at P10 (20). VPS18 rescue cells were generated by transducing clone 1 Hoxb8 cells which still carried the mutation in the *Vps18* gene. Here, the presence of the mutated gene did not affect the maturation when an additional allele of *VPS18* was present during progenitor differentiation to neutrophils. Thus, a certain threshold of VPS18 protein amount might be needed for efficient neutrophil maturation.

Molecular analyses revealed that a heterozygous VPS18 deficiency resulted in the subsequent downregulation of the VPS18 binding partners VPS11, VPS16 and VPS33A. Similarly, homozygous mutations in *VPS16* (c.2272-18C>A) and *VPS33A* (c.1492C>T) were reported to induce the downregulation of CORVET and HOPS core complex components leading to an overall reduction of complex assembly, suggesting that this is a major contributor to disease development (10, 12). This concept is supported by the study from Cai *et al.* in which patients with a different *VPS16* (c.156C>A) mutation did not suffer from neutropenia (30). Tethering complex assembly was not analyzed in *VPS16* (c.156C>A) patients but the above-mentioned phenotype of the *VPS16* (c.2272-18C>A) on CORVET/HOPS subunit levels could be rescued with mutated *VPS16* (c.156C>A) indicating that the *VPS16* (c.156C>A) mutation did not interfere with complex assembly and that disease development of primary dystonia was induced differently (10). Patients with homozygous mutations in *VPS11* (c.2536T>G) are affected by leukoencephalopathy disorder but clinical immune functions seemed not to be affected (31). Mutations in *VPS11* were located in the C-terminally located RING domain which was reported to be nonessential for complex assembly and the authors speculated that HOPS and CORVET complex assembly was unaffected in these patients (32). However, a later study reported that these mutations in *VPS11* led to increased ubiquitination and turnover of VPS11 in HeLa cells and reduced interaction with VPS18 suggesting reduced complex assembly in these cells (33). Overall, CORVET/HOPS complex assembly/stability might be key for the disease phenotype, but cell type specific expression levels of subunits may further influence the phenotype and hence, mutations in one of the VPS subunits might show a stronger effect on a different cell type than mutations in another VPS subunit.

Based on our observations and the current literature, VPS18 deficiency seems to impair vesicle dynamics due to reduced CORVET/HOPS stability. While we have found evidence for an accumulation of autophagosomes, an impaired autophagosomal flux and a reduced autophagosome-lysosome fusion during neutrophil development, others have found additionally an effect on endosomes. In conditional KO mouse brain (Nestin-*Cre*-Vps18^fl/fl^) of postnatal animals (P10), levels of early endosome antigen 1 and RAB7 were increased and autophagosomes and densely stained vesicles accumulated (20). In *vps18* deficient zebrafish larvae, vesicle accumulation and lysosome depletion was observed (22). These differences in phenotypes might be due to the cell types analyzed, different expression levels of the single subunits and whether mutations are present hetero- or homozygously.

The process of autophagy has been linked to proper neutrophil differentiation (24). Induction of autophagy is necessary during early stages of maturation and declines during later stages before a final increase in autophagy occurs in mature neutrophils (4). Abnormally increased autophagy interferes with neutrophil development and results in reduced numbers of mature neutrophils (24). We propose that due to an accumulation of autophagosomes and reduced autophagolysosomal fusion in *Vps18* Hoxb8 mutants, neutrophil progenitors died by premature apoptosis leading to impaired neutrophil maturation.

In the newly identified patient with a heterozygous mutation in *VPS18* neutrophil numbers were increasing with age. Furthermore, the patient’s father carries the same mutation but had normal neutrophil counts at the time of investigation. This suggests that this *VPS18* mutation might induce a transient neutropenia which can improve with age (34). Similar findings were reported from Schürch *et al*. who investigated neutropenia induced by loss of Srp54 in zebrafish. Another possible explanation is the phenomenon of an incomplete penetrance. This has been observed in patients with *VPS33A* mutations: From five patients analyzed, one presented with normal neutrophil counts although all patients harbored the same mutation (12). Similarly, patients with heterozygous *VPS16* loss-of-function mutations suffering from dystonia were also characterized by an incomplete disease penetrance (35).

In conclusion, we found that heterozygous VPS18 deficiency and subsequently an instability of the tethering complexes resulted in delayed neutrophil development and premature apoptosis. We confirmed our findings *in vitro* in the human iPS cell system and *in vivo* in zebrafish larvae. Additionally, we described a patient with a novel heterozygous mutation in *VPS18* suffering from neutropenia and recurrent infections. Our data indicate that proper vesicle dynamics are necessary for neutrophil development and impairment of vesicle dynamics can result in congenital neutropenia.

## Materials and methods

### Cells, cell culture and cell isolation

Human embryonic kidney (HEK) 293T cells (ATCCCRL-11268), HL-60 cells (ATCC CCL-240) and Chinese hamster ovary (CHO) cells were cultured in RPMI 1640 (Sigma-Aldrich) supplemented with 1% penicillin/streptomycin (Sigma-Aldrich) and 10% FCS (Sigma-Aldrich). CHO cells were a kind gift from H. Häcker (University of Utah) and were used to produce SCF-containing supernatant. For differentiation into neutrophil-like cells, HL-60 cells were maintained in RPMI 1640 with 1.3% DMSO for 6 d. Hoxb8 cells were cultured as described before (36). Wild-type Hoxb8 cells were generated using the plasmids *pMSCVneo-ER-Hoxb8* and *pCL-Eco* as described before (37). Plasmids were kindly gifted from from H. Häcker (University of Utah) (23). For differentiation of Hoxb8 progenitor cells in mature neutrophils, cells were washed with PBS (Biowest) and cultured with RPMI 1640 supplemented with 1% penicillin/streptomycin, 10% FCS, 20 ng/ml recombinant murine (rm)G-CSF (PeproTech), and 2% SCF-containing supernatant for 4 days. Cells were kept in a humified atmosphere at 37 °C with 95% O_2_ and 5% CO_2_. Isolation of murine bone marrow and primary peripheral human neutrophils was carried out as described before (38, 39).

### Generation of VPS18 mutant and VPS18 rescue Hoxb8 cell lines

*VPS18* mutant Hoxb8 cells were generated using clustered regularly interspaced short palindromic repeats (CRISPR)/Cas9 technique. Guide (g)RNA 1 (CGCTCGGCCGTCTTGCAGAC) and gRNA 2 (AGGCTAGTGATCCGCTCCGA) were cloned into the lentiCRISPR v2 vector. Next, HEK293T cells were transfected with Vps18 gRNA1 or 2-containing lentiCRISPR v2, psPAX2 and pCMV-VSV-G to generate virus-containing supernatant, which was used for transduction of WT Hoxb8 cells. As control, WT Hoxb8 cells underwent the same procedure and were transduced with virus generated with an empty lentiCRISPR v2, psPAX2 and pCMV-VSV-G. Upon puromycin selection, transduced Hoxb8 cells were subcloned and screened for mutations in the *Vps18* gene. For genotyping of generated *Vps18* mutants, the following primers were used: forward 5’-CCC AGG ACT CCA GTT AAC ACT C-3’ and reverse 5’-GGG CTT AGA TCC TAA ACA CAC G-3’ (gRNA 1), forward 5’-AGG GGA GCC TAT AAT CTC TTG G-3’ and reverse 5’-TCA AAC TTC GAC CAA TGA CAA G-3’ (gRNA 2). LentiCRISPR v2 was a gift from Feng Zhang (Addgene plasmid # 52961; http://n2t.net/addgene:52961; RRID:Addgene_52961) (40). psPAX2 was a gift from Didier Trono (Addgene plasmid # 12260; http://n2t.net/addgene:12260; RRID:Addgene_12260). pCMV-VSV-G was a gift from Robert Weinberg (Addgene plasmid # 8454; http://n2t.net/addgene:8454; RRID:Addgene_8454) (41). The pEGFP-N1 plasmid containing full-length human (h)*VPS18* was a kind gift of Stephen C. Graham (University of Cambridge) (19). The pMSCV-Puro vector was provided by Hans Häcker (University of Utah). For generation of rescue cells, the coding region of hVPS18 was cloned into the pMSCV-Puro vector. Stable transduction of Hoxb8 cells was carried out as described before (36). In brief, HEK293T cells were transfected with pMSCV-Puro-hVPS18-EGFP and pCL-Eco using Lipofectamine 2000 and virus-containing supernatant was collected. Clone 1 Hoxb8 cells were transduced by spinoculation using Lipofectamine. Puromycin selection was started 72 h post transduction for 7 days. Subsequently, rescue cells were sorted for their EGFP expression. VPS18-EGFP fusion protein expression was analyzed by flow cytometry and Western blot.

### Cytospins and May-Grünwald-Giemsa staining

Cytospins were generated with a Cellspin III Cytocentrifuge (THARMAC). 5×10^4^ Hoxb8 cells were centrifuged at 1,000 rpm for 10 min onto glass slides. For May-Grünwald-Giemsa staining, cytospins were stained 3 min with May-Grünwald solution (Merck Millipore) followed by staining with 1.3% Giemsa solution (Merck Millipore) for 20 min. Images were acquired using a 100x/1.40 Oil objective of the Leica DM2500 microscope equipped with a DMC2900 CMOS camera (Leica).

### Immunoblotting

Whole cell lysates were generated as described before (42). The following specific primary and appropriate secondary antibodies were used: anti-VPS18 (abcam, EPR13378), anti-LC3B (Cell Signaling Technology, #2775), anti-RAB5 (LS Bio, LS-B12415-300), anti-RAB7 (abcam, EPR7589), anti-GFP (Proteintech, 1E10H7), anti-ATF6 (Proteintech, 24169-1-AP), anti-GAPDH (Merck Millipore, 6C5), anti-beta Tubulin (Sigma Aldrich, 2-28-33), anti-beta Actin (Santa Cruz Biotechnology, C4) and IRDye 800CW-labeled anti-rabbit (LI-COR Biosciences, #926-32213), IRDye 680RD-labeled anti-goat (#925-68074), IRDye 680RD-labeled anti-mouse (#926-68072). Immunoblots were imaged with the Odyssey CLx (LI-COR Biosciences) and analyzed with Image Studio Lite (LI-COR Biosciences).

### Flow cytometry of Hoxb8 cells

For flow cytometry, the following antibodies were used: anti-VPS18 (abcam, EPR13378), Alexa Fluor (AF)488-labeled secondary antibody (Invitrogen), AF488-labeled anti-p62 antibody (abcam, EPR4844), PE-conjugated anti-active caspase 3 antibody (BD, C92-605), anti-EIF2S1 (phospho S51, abcam, 10H21L20), anti-eIF2α (abcam, EPR11042), anti-XBP-1s (Cell Signaling Technology, E9V3E) labeled with AF488 using the FlexAble Antibody Labeling Kit (Proteintech), AF647-labeled anti-CHOP antibody (Biolegend, 9C8) and the appropriate isotype controls. For intracellular stainings, Hoxb8 cells were fixed in fixation/permeabilization solution (BD Biosciences) and incubated with the appropriate primary antibody or isotype control antibody in BD Perm/Wash buffer (BD Biosciences) at 4°C for 45 min. In case of unlabeled primary antibodies, secondary antibody was subsequently applied at 4 °C for 45 min. For cell surface stainings, Hoxb8 cells were incubated with the desired antibodies for 20 min at 4 °C and fixed using Fixation Buffer (Biolegend) for 15 minutes at RT. Dead cells and doublets were excluded using Zombie Aqua (Biolegend), LIVE/DEAD^TM^ Fixable Yellow or Near-IR Dead Cell Stain (Invitrogen). For staining of lipid droplets, Hoxb8 cells were incubated with 10 µg/ml Nile red (Thermo Fisher Scientific) or 2 µM BODIPY™ 493/503 (Thermo Fisher Scientific) for 15 min at 37 °C in PBS. For staining of acidic compartments, cells were stained with 50 nM LysoTracker^TM^ Red DND-99 (Thermo Fisher Scientific) for 45 min at 37 °C in 5% FCS in PBS. Dead cells were excluded using LIVE/DEAD^TM^ Fixable Near-IR dead cell stain or SYTOX Red dead cell stain (Invitrogen, #S34859). For cell death analysis, cells were stained using FITC-Annexin V (Biolegend, #640906), SYTOX Red dead cell stain and tetramethylrhodamine methyl ester (TMRM, Invitrogen, #T668). If not stated otherwise, Hoxb8 cells were analyzed using a CytoFLEX S flow cytometer (Beckman Coulter). Data were analyzed with FlowJo^TM^ software (BD Biosciences).

### Spectral flow cytometry of Hoxb8 cells

For analysis of neutrophil maturation via spectral flow cytometry, cells were incubated with the following antibodies: BUV395-labeled anti-Ly6G (BD, 1A8), BUV496-labeled anti-CD11b (BD, M1/70), BUV615-labeled anti-CD16/32 (BD, Ab93), BUV661-labeled anti-Sca-1 (BD, D7), BUV563-labeled anti-CD49d (BD, 9C10), BUV737-labeled anti-CD115 (BD, AFS98), BV421-labeled anti-c-kit (Biolegend, 2B8), BV480-labeled anti-CD62L (BD, MEL-14,), BV605-labeled anti-CD106 (BD, 429), BV711-labeled anti-CXCR4 (Biolegend, L276F12), BV785-labeled anti-Ly6C (Biolegend, HK1.4), PerCP/Cy5.5-labeled anti-CD81 (Biolegend, Eat-2), AF647-labeled anti-CD34 (Biolegend, HM34), APC-labeled anti-CD101 (Invitrogen, Moushi101), PE-labeled anti-CXCR2 (Biolegend, SA044G4) and AF700-labeled anti-CD24 (Biolegend, M1/69). Incubation was carried out for 20 minutes at 4 °C and fixed using Fixation Buffer (Biolegend) for 15 minutes at room temperature. Dead cells and doublets were excluded using forward and side scatter characteristics and auto-flourescence signal. Data acquisition was performed on a Cytek® Aurora (Cytek Biosciences) and data were analyzed using FlowJo v10.10.0 (BD Bioscience).

### Transmission electron microscopy

Samples were fixed with 2.5% glutaraldehyde in 0.1 M sodium cacodylate buffer, pH 7.4 (Electron Microscopy Sciences) for 24 h at minimum. Thereafter, glutaraldehyde was removed and samples were washed with 0.1 M sodium cacodylate buffer, pH 7.4. Postfixation and prestaining was done for 45 to 60 min with 1% osmium tetroxide (Electron Microscopy Sciences), Samples were washed with ddH_2_O and dehydrated with an ascending ethanol series (15min with 30%, 50%, 70%, and 90% respectively and two times 10 min with 100%). Subsequently, samples were embedded in Epon (Serva Electrophoresis GmbH). 60-70 nm thick ultrathin sections were cut at the Reichard-Jung Ultracut E microtome. Ultrathin sections were collected on formvar coated copper grids (Plano) and automatically stained with UranyLess EM Stain (Electron Microscopy Sciences) and 3% lead citrate (Leica) using the contrasting system Leica EM AC20 (Leica). Imaging was carried out using the JEOL -1200 EXII transmission electron microscope (JEOL) at 80 kV. Images were taken using a digital camera (KeenViewII; Olympus) and processed with the iTEM software package (anlySISFive; Olympus).

### MS-based proteomics and MS data analysis

Cell pellets were resuspended in 50 µL digestion buffer (1% sodium deoxycholate, 40 mM tris(2-carboxyethyl)phosphine, 40 mM 2-chloroacetamide in 100 mM Tris, pH 8.5) and heated for 10 min at 95 °C, 1400 rpm. Samples were sonicated for 15 min at 4 °C and digested with 1 µg LysC/trypsin mix for 18 h at 37 °C, 1000 rpm. Digestion was stopped by adding 300 µL isopropanol, 1% trifluoroacetic acid (TFA). Peptides were desalted using in-house made SDB-RPS StageTIPS and resuspended in 12 µL of buffer A* (2% acetonitrile, 0.1% TFA) for LC-MS. Desalted peptide mixtures were analyzed with an EASY-nLC 1200 HPLC system (Thermo Fisher Scientific) coupled to an Orbitrap Exploris 480 instrument (Thermo Fisher Scientific) via a nano-electrospray ion source. 250 ng peptides were loaded onto an in-house packed column (75 μm inner diameter, 50 cm length, packed with 1.9 μm C18 ReproSil beads (Dr. Maisch GmbH)). To elute peptides, a buffer system was used consisting of buffer A (0.5% formic acid) and buffer B (0.5% formic acid, 80% acetonitrile) with a linear gradient from 5% to 30% buffer B in 120 minutes at flow rate of 300 nl/min. The column temperature was maintained at 60°C. MS data were acquired using the data-independent acquisition (DIA) mode operated by the XCalibur software (Thermo Fisher). DIA was performed with one full MS survey scan followed by 33 MS/MS windows in one cycle. Survey scans were acquired at 120,000 resolution with an AGC target of 3×10^6^ charges in the 300–1650 m/z range and a maximum injection time of 60 ms. DIA precursor windows ranged from 300 m/z (lower boundary of first window) to 1,650 m/z (upper boundary of last window). MS/MS settings included a resolution of 30,000 at m/z 200, an AGC target of 3×10^6^ charges and a maximum injection time of 54 ms. MS raw files were processed using Spectronaut (version 14.3.200701.47784) and mass spectra were searched against the mouse UniProt FASTA database (July 2019, 63,439 entries) with an FDR of 1% at the protein and peptide level. A minimum peptide length of 7 amino acids and a maximum of two missed cleavages were allowed in the database search. Protein groups were filtered such that only proteins identified in at least 3 replicates of one condition were retained. Missing values were replaced from a Gaussian distribution (30% width and downshift by 1.8 standard deviations of measured values) and t-tests were applied with a permutation-based FDR of 5%. Data and statistical analysis were done in R (v4.0.3).

### Maintenance and differentiation of human iPSCs

The human fibroblast-derived iPS cell line HMGU1 (iPSC1) was provided by Micha Drukker at the iPSC Core Facility, Institute of Stem Cell Research, Helmholtz Center Munich (hPSCreg IFSi001-A) and used as healthy control cell line. Authentication and assessment of genomic integrity were performed by the provider. The *VPS18* mutant derivative cell lines are described below. iPSCs were maintained on a tissue culture dish coated with growth factor-reduced Matrigel (Corning) in mTeSRplus serum-free medium (Stemcell Technologies) at 37 °C and 5% CO_2_. Cells were passaged every 3-4 days using ReLeSR (Stemcell Technologies). Differentiation of iPSCs into neutrophil-like cells was carried out as described before. (28) For determination of live floating cells during neutrophil-directed differentiation, the number of viable cells in suspension (live floating cells representing all cellular subsets of neutrophil-directed differentiation) was determined at day 18, 21, 25 and 28 of differentiation by the Trypan Blue (Gibco) staining assay.

### CRISPR/Cas9-mediated genome engineering of iPSCs

For the generation of *VPS18*-deficient iPS cells, the targeting sequence AGACTGGTAATGCGCTCGGA (gRNA1) of the *VPS18* gene was selected. For generation of iPSCs with the patient-specific genetic variant Arg234* the CTCGGCCTATGAACTGGAAG (gRNA2) was used. gRNAs were inserted into the PX458 plasmid (pSpCas9(BB)-2A-GFP (PX458) was a gift from Feng Zhang (Addgene plasmid # 48138 ; http://n2t.net/addgene:48138 ; RRID:Addgene_48138)). 10^6^ iPSCs were resuspended with supplement and nucleofector solution according to the human stem cell nucleofector Kit 2 (Lonza) followed by the addition of 5 μg of the plasmid encoding the gRNA targeting *VPS18*. For generation of the knock-in Arg234*, 10^6^ iPSCs were co-transfected with 5 μg of the plasmid encoding the gRNA targeting VPS18 and 160 pmol of the donor template GAGCGGGGCCCTGATGGGCGTAGCTTTGTTATTGCCACCACTCGGCAGCGCCTCTTCCAGTTCATAGGCtGAGCAGCAGAGGGGGCTGAGGCCCAGGGTTTCTCAGGGCTCTTTGCAGCTTACACGGACCACCCACCCCC (ssODN, Integrated DNA Technologies). Upon resuspension, cell suspensions were transferred into a cuvette and nucleofected by the Amaxa Nucleofector® II Device (Lonza, program B-016). iPSCs were then transferred to a tissue culture dish coated with growth factor-reduced Matrigel (Corning) in mTeSR1 serum-free medium (Stemcell Technologies) at 37 °C and 5% CO_2_. After 8 h, SCR7 (10 μM) (Stemcell Technologies) was added and removed after 12 h. Two days after transfection, GFP-positive iPSCs were sorted by BD FACS Aria (BD Biosciences) and 3000 sorted iPSCs were seeded on a 10 cm dish coated with growth factor-reduced Matrigel in mTeSR1 serum-free medium. Around 10 days after seeding, each colony derived from a single cell was picked up for genotyping and further expansion. Genomic DNA was PCR-amplified using OneTaq Polymerase (NEB). The following primers were used for genotyping: 5’-TCACCTGTCTCTTCCACAGCTA-3’ (forward, VPS18 KO), 5’-TTAACAGTCCCCAAACACCTCT-3’ (reverse, VPS18 KO) and 5’-GCTACAGTGAGTTGGCCTTCTA-3’ (forward, VPS18 Arg234*), 5’-AGTAGCAGCAGGAAGTGGAACT-3’ (reverse, VPS18 Arg234*). Amplicons were sequenced at Eurofins Genomics.

### Immunoblotting and flow cytometry of iPSCs

For immunoblot analyses, iPSCs were lysed in Laemmli buffer, cell lysates were subjected to SDS-PAGE and transferred to a nitrocellulose membrane. Membranes were blocked for 1 h at room temperature in 5% non-fat milk. All antibodies were diluted in 5% non-fat milk. The following primary antibodies were used: anti-LC3B (Cell Signaling, #2775), anti-VPS18 (abcam, EPR13378), anti-beta actin (Santa Cruz, C4), anti-VPS11 (Proteintech, #19140-1-AP), anti-VPS16 (Proteintech, #17776-1-AP) and anti-VPS33a (Proteintech, #16896-1-AP). Horseradish-peroxidase-conjugated anti-rabbit (Cell Signaling, #7074S) was used as secondary antibody.

Flow cytometry analyses were performed as previously described (29). Briefly, iPSCs were harvested at day 28 of differentiation and washed once with FACS buffer (2% FBS in PBS). iPSCs were resuspended in 50 μl FACS buffer supplemented with human Trustain FcX (Biolegend, cat# 422302) for 7 min at RT and then stained with the following antibodies or fluorescent stains in Brilliant stain buffer (BD, cat# 563794) for 20 min at RT in the dark: Fixable Viability Stain 780 (BD, cat# 565388), PE anti-CD49d (BD, cat#555503), BB515 anti-human CD11b (BD, cat# 564518), PE-Cy7 anti-human CD16 (BD, cat#557744), APC-R700 anti-human CD45 (BD, cat# 566041), BV480 anti-human CD35 (BD, cat# 746503), BV605 anti-human CD101 (BD, cat# 747548), APC anti-human Siglec-8 (BioLegend, cat# 347106), APC/Fire750 anti-human CD14 (BioLegend, cat# 301854). Afterwards, cells were washed with FACS buffer and resuspended in Annexin binding buffer. Cells were stained with Pacific Blue Annexin V (Biolegend, cat# 640918) according to the instructions by the manufacturer. Cells were then strained through the mesh of a round bottom polystyrene test tube (BD, cat# 352235) and subsequently analyzed by the BD FACS Aria (BD Biosciences).

### Zebrafish husbandry, genetic manipulation and tail fin transection

Zebrafish larvae were kept at 28.5 °C in E3 medium (5 mM NaCl, 0.33 mM CaCl2, 0.33 mM MgSO4, 0.17 mM KCl, 0.00003% methylene blue). At 24 hpf, 1-Phenyl 2-thiourea (0.003%, Sigma-Aldrich) was added to the medium. Adult zebrafish were raised and housed as well as the experiments in zebrafish larvae were performed in accordance with animal protection standards of the Ludwig-Maximilians-Universität München and approved by the government of Upper Bavaria (Regierung von Oberbayern). In this study, zebrafish embryos of the transgenic line Tg*(fli1:gfp;lyz:dsred)* were analyzed and used to generate the two *vps18*-mutant lines Tg*(fli1:gfp;lyz:dsred;vps18mde403)* and Tg*(fli1:gfp;lyz:dsred;vps18mde404)* referred to as Vps18 wild-type (WT), Vps18 mutant 1, and Vps18 mutant 2, respectively (43–45). For genome editing using CRISPR/Cas9 technique, gRNAs (IDT) targeting exon 1 (gRNA 1, CGTATGTCCACGGCCAACAT) and exon 4 (gRNA2, CATTGAGGCCAAGCAAGAGG) were designed. Cas9 protein (IDT) and gRNAs were injected into zebrafish embryos at one cell stage as described before (43). Adult *vps18* founders were outcrossed with Tg*(fli1:gfp;lyz:dsred)* zebrafish and offspring was genotyped for germ line mutations in *vps18*. Genotyping was performed with the following primers: 5’-AAGACAGACATGCAACCAACAC-3’ (forward), 5’-AATGGGTTTTTCTTCCTCCAAT-3’ (reverse) (gRNA1) and 5’-TAGTCCTCACCCAGTTCCACTT-3’ (forward), 5’-CCGATTCAGATAGAGTTCGGTC-3’ (reverse) (gRNA2). PCR products were sequenced to verify amplification of correct DNA sequences. Neutrophil trafficking during sterile inflammation was analyzed following tail fin transection as described before (43).

### Total neutrophil count in zebrafish larvae

Total neutrophil numbers were analyzed in zebrafish larvae 3 dpf upon euthanization with an overdose of tricaine (0.3 mg/ml), fixation in 4% paraformaldehyde and washing with PBS. Larvae were imaged with an upright spinning disc confocal laser microscope (Examiner, Zeiss) equipped with a confocal scanner unit CSU-X1 (Yokogawa Electric Corporation), a CCD camera (Evolve, Photometrics) and a 5x/0.15NA objective (N-Achroplan, Zeiss) and Slidebook 6.0.13 Software (3i). Quantification was carried out by manual counting in a blinded fashion using the Cell Counter plugin in FIJI (46).

### Statistics

If not stated otherwise, GraphPad Prism was used for statistical analysis. All datasets were analyzed with parametric tests (Student’s *t* test, one- or two-way ANOVA) followed by post hoc multiple comparisons. *P* values were considered significant with \**P*<0.05, \*\**P*<0.01, \*\*\**P*<0.001, and \*\*\*\**P*<0.0001.

## Supporting information

Gao et al_Supplementary Material

## Acknowledgments

The authors thank Jennifer Truong, Tanja Vlaovic, Tanja Weißer, Ulrike Wilhelm-Forster, Sabine Schlink and Hafez Gabara for excellent technical assistance. The authors are grateful to Dr. Steffen Dietzel (Core Facility Bioimaging, Biomedical Center, LMU Munich) for the support with fluorescence microscopy. We acknowledge the Core Facility Flow Cytometry at the Biomedical Center, LMU Munich, for providing equipment and expertise.

## Author contributions

D.M.-B. designed and performed experiments, analyzed data and wrote the manuscript. J.G., A.B., M.I.L., J.C., F.M., M.Ri., A.Z., K.M., M.Ro., M.T., B.P. and O.S. performed experiments and analyzed data. S.F.-W., B.S., C.K. and B.W. provided their expertise. I.S., J. Y., O.S.-S. and R.S. provided critical patient information. D.M.-B. and B.W. acquired funding and supervised the study.

