## Supplementary material for "Mutations in *VPS18* lead to a neutrophil maturation defect associated with disturbed vesicle homeostasis": Gao et al_Supplementary Material

### Suppelementary methods

#### PCR

Gene expression analysis was carried out as described before (1). In brief, RNA isolation was carried out using the RNeasy Mini Kit (QIAGEN) according to the manufacturer's protocol. cDNA synthesis was conducted with the Maxima First Strand cDNA Synthesis Kit (Thermo Fisher Scientific). For VPS18 gene expression analysis, the following primers were used: 5'-CCTGCAGGTGGATGTGGACC-3' (forward, murine), 5'-GCGCTGCAGTTCCTCAAGTC-3' (reverse, murine), 5'-CCTGCAGGTGGATGTGGACC-3' (forward, human), 5'-GCAGGCAGTCAGCATGGAAC-3' (reverse, human). For GAPDH expression as loading control the following primers were used: 5'-GGGCTCATGACCACAGTCCA-3' (forward, murine), 5'-GAGGTCCACCACCCTGTTGC-3' (reverse, murine), 5'-GGGGAGCCAAAAGGGTCATCATCT -3' (forward, human), 5'-TGTGCTCTTGCTGGGGCTGGTG -3' (reverse, human). Genomic (g)DNA for genotyping was isolated using PCRBIO Rapid Extract PCR Kit (PCR Biosystems). For amplification of cDNA or gDNA, PCRBIO HS Taq Mix (PCR Biosystems) with appropriate primer pairs were used in a peqSTAR thermocycler (peqlab). PCR products were separated in 2% agarose gels and stained with MIDORI Green (NIPPON Genetics). Amplification of correct DNA sequences was verified by sequencing of the PCR products.

#### Immunofluorescence stainings, image acquisition and analysis

For the analysis of subcellular localization of Rab5, Rab7 and LAMP1, the following antibodies were used: Cy3-conjugated anti-LAMP1 (abcam, ab67283), AF647-conjugated anti-Rab7 (abcam, EPR7589), anti-Rab5 (LS Bio, LS-B12415-300), AF488-labeled anti-goat (LS Bio, LS-C149359). dHoxb8 cells (1.5 x $10^5$ cells/well) were immobilized on rmICAM-1 (3 µg/ml, Stemcell) and rmCXCL1 (5 µg/ml, R&D Systems) in wells of a 12-well chamber (Ibidi), fixed in 4% PFA (Sigma-Aldrich) and permeabilized with 0.5% saponin (Sigma-Aldrich) and 10% bovine serum albumin (BSA, Sigma-Aldrich). Upon blocking in 10% BSA, primary antibodies were incubated overnight at 4 °C and secondary antibodies for an hour at RT in 0.5% saponin and 10% BSA. The nucleus was stained with Hoechst 33342 (Thermo Fisher Scientific). Analysis of subcellular localization of Rab5, Rab7, and LAMP1 was performed using an inverted Leica SP8X white light laser microscope and an 100x/1.4-NA oil immersion objective (Leica). The images were deconvoluted using Huygens Deconvolution software (Scientific Volume Imaging B.V.) prior to image analysis. Images were analyzed with Leica Application Suite X 3.4.2.18368 software (Leica) and ImageJ (NIH) (2). Z-stack images of at least four representative cells were taken for each immunofluorescence staining. The images are presented as a maximum projection or a single z-stack layer.

**Analysis of whole-kidney marrow in adult zebrafish**

Adult zebrafish were euthanized at two years of age using 0.3 mg/ml tricaine in E3 medium. Isolation of the kidney was carried out as described by others (3). To generate a single cell solution, the organ was gently pressed over a 40 µm cell strainer using a plunger of a 1ml syringe. SYTOX Red dead cell stain was used to label and exclude dead cells. Analysis of whole-kidney marrow (WKM) cells was conducted with the CytoFLEX S flow cytometer (Beckman Coulter) and data were analyzed with FlowJo™ software (BD Biosciences) as described by Traver et al. (4). DsRed-positive cells were identified as neutrophils.

**Table S1. Serial absolute neutrophil count (ANC) from patient 1.**

| Age | ANC (K/µl) |
|---|---|
| 9 mo | 0.3 |
| 10 mo | 0.4 |
| 12 mo | 0.2 |
| 1.2 y | 0.7 |
| 1.5 y | 1.0 |
| 1.9 y | 0.6 |
| 1.10 y | 1.4 |
| 1.11 y | 8.7 (acute infection) |
| 2 y | 0.3 |
| 2.3 y | 0.7 |
| 2.5 y | 1.7 |
| 2.7 y | 0.9 |
| 2.10 y | 2.2 |
| 3.2 y | 3.3 |
| 3.7 y | 3.4 |
| 4.11 y | 2.4 |

**Table S2. Detailed patient information.**

| Patient-ID | Gene | Ensemble transcript ID | HGVSC | Genomic position | CADD v1.6 | gnomAD AF (V4.0.0) | gnomAD count | Zygosity | Effect on protein | HGVSP |
|---|---|---|---|---|---|---|---|---|---|---|
| P1 | *VPS18* | ENST00000220509 | c.700C>T | 41191716 | 27,3 | 2.74e-6 | 4 | Het | stop-gained | p.Arg234Ter |

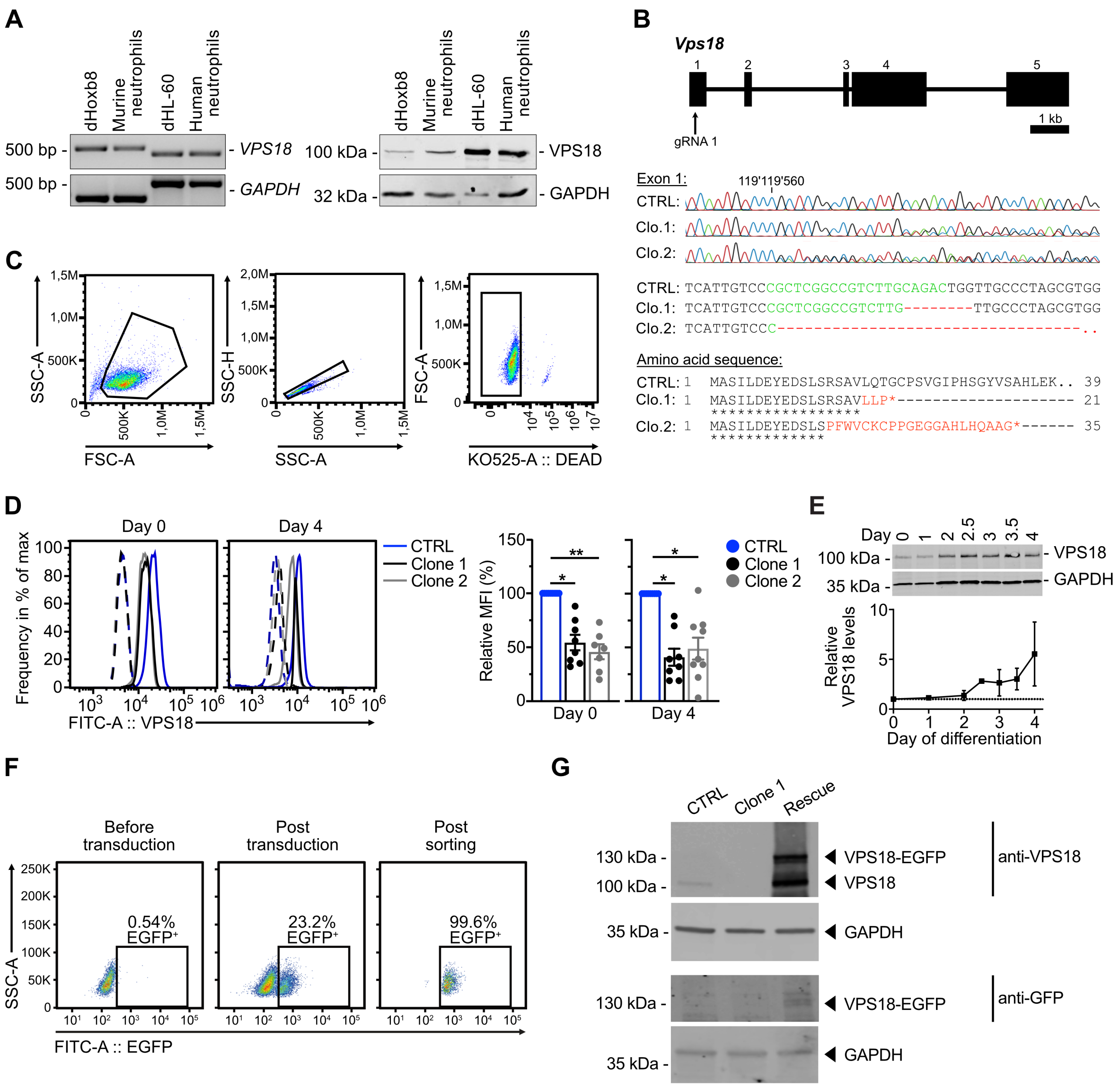


**Supplementary Fig. 1. Generation of *Vps18* mutant Hoxb8 cell lines and gating strategies of VPS18 detection.** (A) Representative images of VPS18 expression on mRNA (left panel) and protein level (right panel) in murine dHoxb8 and human dHL-60 cells as well as freshly isolated murine bone marrow and human peripheral blood neutrophils. mRNA was detected using RT-PCR, protein was detected by Western blot technique. GAPDH was used as loading control. n = 3. (B) Upper panel: Schematic of murine *Vps18* gene and targeted exon 1 of gRNA 1. Middle panel: Partial genomic sequences of control and *Vps18* mutant Hoxb8 cell lines (CTRL, clone (clo.) 1 and 2). Green, guide (g)RNA sequence. Red, deletions. Numbers indicate position within the chromosome. Lower panel: Predicted amino acid sequence of mutants aligned to CTRL sequence of the first 39 amino acids. Identical (*) and altered (red) amino acids are indicated. (C) Gating strategy for single, living cells of Hoxb8 cells from CTRL, clone 1 and clone 2 Hoxb8 cell lines. Representative gating strategy for CTRL is shown. (D)

Representative histograms (left panel) and quantification (right panel) of residual VPS18 expression before (day 0) and after removal (day 4) of estrogen in CTRL, clone 1 and clone 2 Hoxb8 cells analyzed by flow cytometry using specific antibodies. Isotype controls, dashed lines. n≥7 for each cell line. Mean ± SEM. *$P < 0.05$, **$P < 0.01$ compared to CTRL. One-way ANOVA, Tukey's multiple comparisons test. (E) Representative Western blot of VPS18 expression (upper panel) and quantitative analysis (lower panel) in cell lysates from CTRL Hoxb8 cells during differentiation (day 0-4). Ratio of VPS18/GAPDH was calculated and presented as relative protein levels. n ≥ 3. (F) Detection of $EGFP^{+}$ cells using flow cytometry in VPS18 rescue cells, before (left panel) and after transduction with *pMSCV-Puro-hVPS18-EGFP* (middle panel) and after FACS (right panel). Numbers represent cells in percent of total cells analyzed per panel (100%). (G) Representative Western blot indicating the expression of VPS18-EGFP and endogenous VPS18 in transduced VPS18 rescue Hoxb8 cells at day 0 of differentiation using specific antibodies. GAPDH was used as loading control.

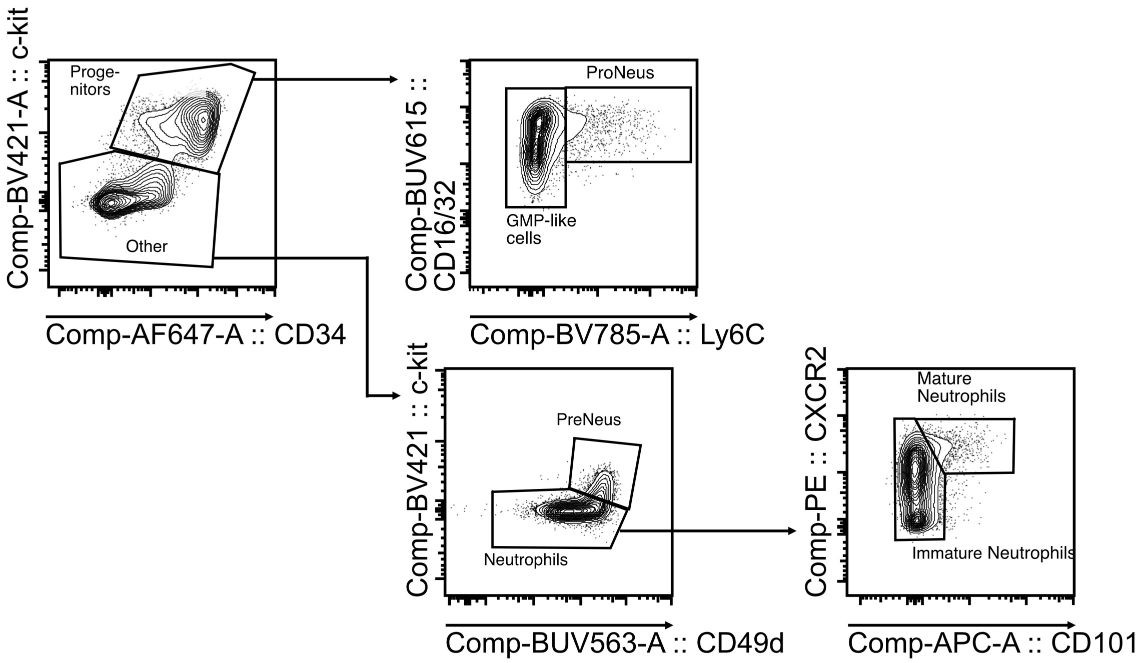


**Supplementary Fig. 2. Gating strategy of neutrophil differentiation based on cell surface markers.** Gating strategy for spectral flow cytometric analysis followed by dimensional reduction using UMAP for $CD34^{hi}$c-$kit^{hi}$ $CD16/32^{+}Ly6C^{-}$ (GMP-like cells), $CD34^{+}$c-$kit^{+}CD16/32^{+}Ly6C^{+}$ (proNeu), $CD34^{-}$c-$kit^{int}CD49d^{+}$ (preNeu), c-$kit^{-}CD49d^{-}CXCR2^{+}CD101^{-}$ (immature Neu) and c-$kit^{-}CD49d^{-}CXCR2^{+}CD101^{+}$ (mature Neu) in all single, living $CD115^{-}$ CTRL, clone 1 and clone 2 Hoxb8 cells. Representative gating strategy for CTRL is shown.

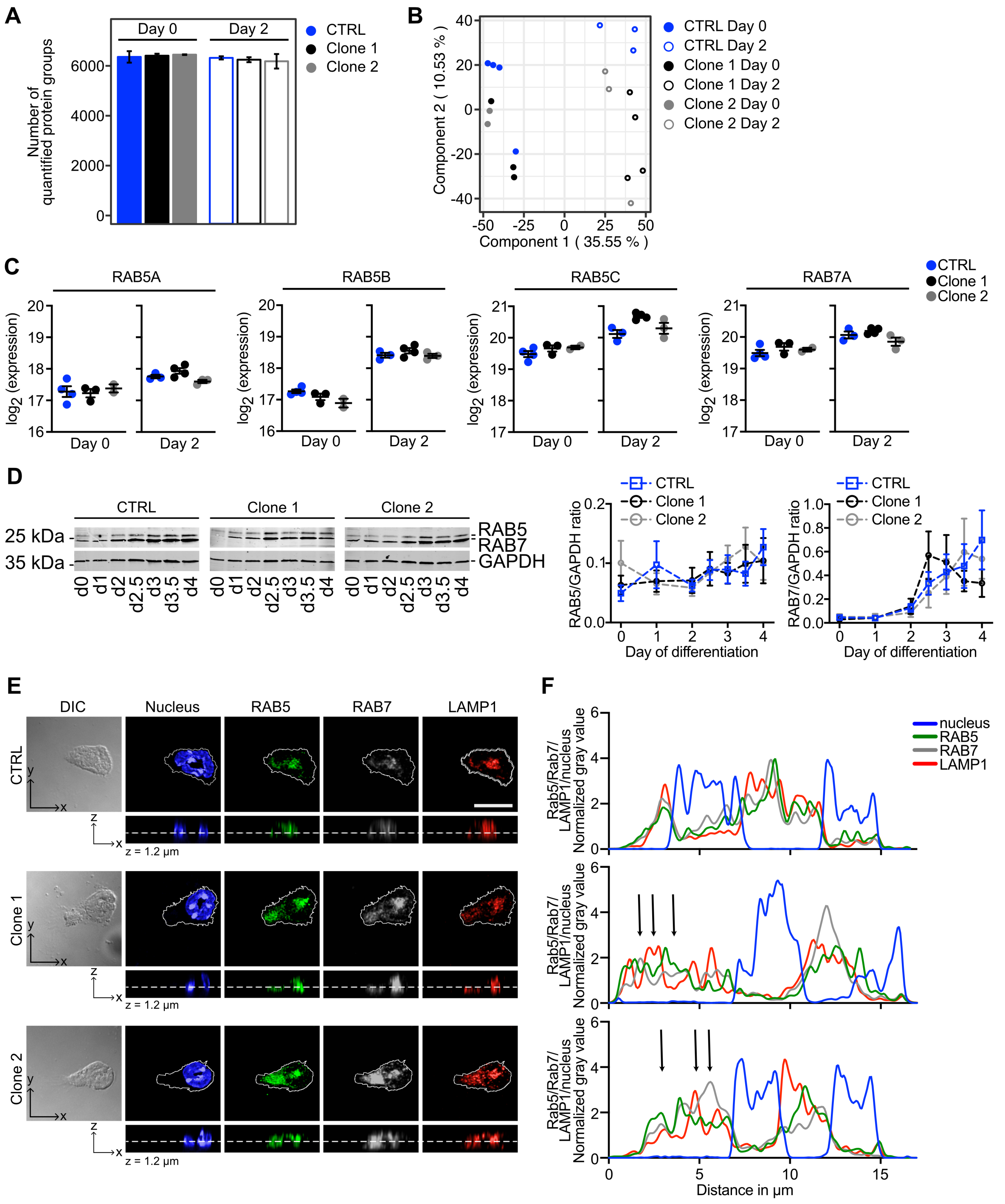


**Supplementary Fig. 3. Disturbed intracellular vesicle homeostasis in *Vps18*$^{+/-}$ mutant Hoxb8 cells.** (A) Number of MS/MS-based quantified protein groups in indicated cell lines at day 0 and 2 of differentiation. Mean ± SEM. n ≥ 2. (B) Principal component analysis for CTRL, clone 1 and clone 2 Hoxb8 cells before (day 0) and during (day 2) differentiation. (C) Expression levels of endosomal markers RAB5A-C and RAB7A in indicated cell lines before (day 0) and during (day 2) differentiation analyzed by mass spectrometry. n ≥ 2. Mean ± SEM.

Pairwise t-test, multiple hypothesis correction. (D) Representative Western blot (left panel) and quantitative analysis (right panel) of RAB5 and RAB7 expression in cell lysates from indicated cell lines during differentiation (day 0-4). Ratios of RAB5/GAPDH and RAB7/GAPDH were calculated and presented as relative protein amount. $n = 7$. Mean ± SEM. Two-way ANOVA, Tukey's multiple comparisons test. (E) Representative images of CTRL, clone 1 and clone 2 dHoxb8 cells upon recombinant murine (rm)CXCL1-induced adhesion to rmICAM-1. Nucleus (blue), RAB5 (green), RAB7 (grey), LAMP1 (red) in one z-stack (1.2 µm) position and in one orthogonal layer. Scale bar, 10 µm. $n = 4$. (F) Exemplary intensity plot profiles of RAB5, RAB7, LAMP1 and nucleus in 1.2 µm z-stack position in indicated cell lines from (B). Arrows indicate shifted localization of analyzed proteins.

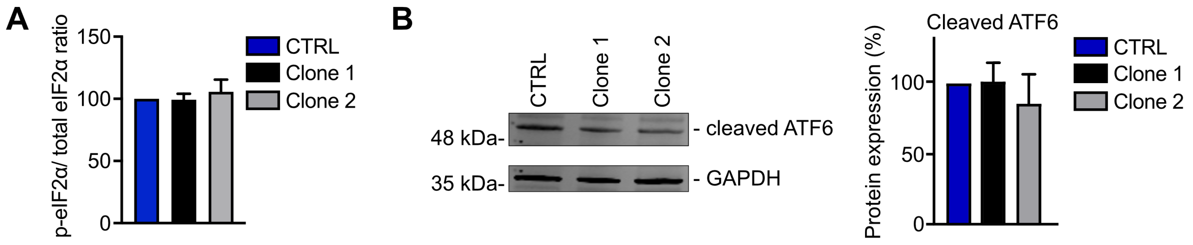


**Supplementary Fig. 4. *$Vps18^{+/-}$* mutant neutrophil progenitors experience cell stress during differentiation.** (A) Levels of phosphorylated eIF2α (p-eIF2α) in indicated cell lines at day 3 of differentiation presented as ratio of p-eIF2α to total eIF2α analyzed by flow cytometry and normalized to CTRL. n = 3. (B) Representative Western blot (left panel) and quantitative analysis (right panel) of cleaved ATF6 in cell lysates from indicated cell lines at day 3 of differentiation. Protein expression was normalized to GAPDH. Ratios of cleaved ATF6/GAPDH were calculated, presented as protein expression normalized to CTRL (100%). n = 3. (A-B) Mean ± SEM. One-way ANOVA, Tukey's multiple comparisons test.

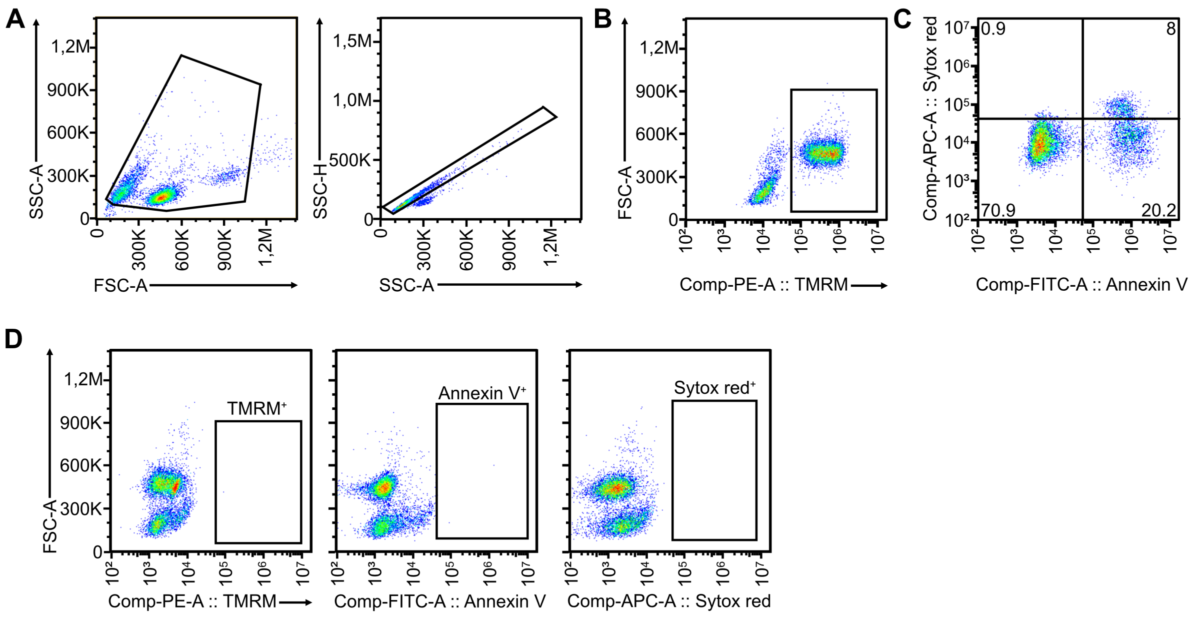


**Supplementary Fig. 5. *Vps18*$^{+/-}$ mutant neutrophil progenitors undergo premature apoptosis.** (A-C) Gating strategy in single cells (A) of CTRL, clone 1 and clone 2 Hoxb8 cell lines for detection of TMRM$^+$ (B), Annexin V$^+$ and Sytox Red$^+$ cells (C). Representative gating strategy for CTRL Hoxb8 cells at day 4 is shown. (D) Fluorescence minus one-controls for TMRM$^+$, Annexin V$^+$ and Sytox Red$^+$ samples.

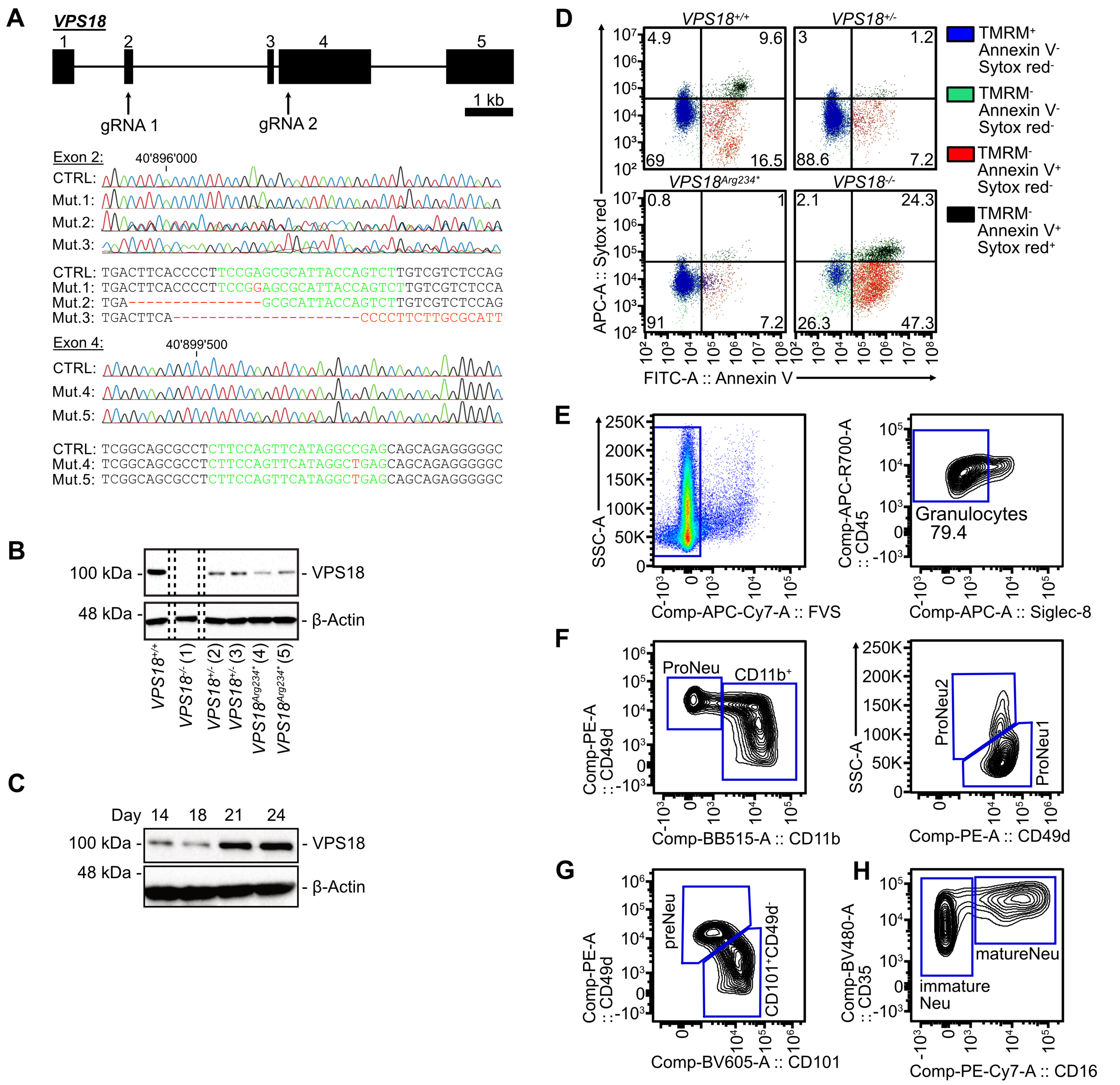


**Supplementary Fig. 6. *VPS18* mutant human iPSC-derived neutrophil progenitors show defects in neutrophil development.** (A) Upper panel: Schematic of the human *VPS18* gene and target exons 2 and 4 of the gRNA1 (for KO mutants) and 2 (for KI mutants). Lower panel: Sequencing traces and partial genomic sequence of *VPS18*$^{+/+}$ cells (CTRL), *VPS18*$^{-/-}$ mutant 1 (mut. 1), *VPS18*$^{+/-}$ mut. 2 and 3, and patient specific *VPS18*$^{Arg234*}$ mut. 4 and 5. Numbers indicate position within the gene. Target sequence of the gRNAs in green. Nucleotides mutated or deleted in red. (B) Representative Western blot of VPS18 expression in cell lysates of iPSC-derived neutrophil progenitors (day 28) in indicated cell lines. β-actin was used as loading control. n = 3. (C) Representative Western blot of VPS18 expression during neutrophil development in cell lysates of wild-type iPSC-derived neutrophil progenitors. β-actin was used as loading control. n ≥ 2. (D) Representative dot plots of viable, preapoptotic, early and late apoptotic cells of indicated genotypes during differentiation (day 28). Viable cells, TMRM$^{+}$, blue. Preapoptotic cells, TMRM$^{-}$, Annexin V$^{-}$ and SytoxRed$^{-}$, green. Early apoptotic cells, Annexin V$^{+}$ and SytoxRed$^{-}$, red. Late apoptotic cells, Annexin V$^{+}$ and SytoxRed$^{+}$, black. Numbers indicate % of cells of all single cells. n ≥ 1. (E-H) Gating strategy for characterization of neutrophil maturation states during differentiation of iPSC out of all single cells. Living, neutrophil-lineage directed cells (E), proNeu1 (CD11b$^{-}$, CD49d$^{+}$, SSC-A$^{low}$) and proNeu2 (CD11b$^{-}$, CD49d$^{+}$, SSC-A$^{high}$) (F), preNeu (CD49d$^{+}$, CD101$^{-}$) (G) and immature (CD35$^{+}$, CD16$^{-}$) and mature (CD35$^{+}$, CD16$^{+}$) neutrophils (H). Representative gating strategy for VPS18$^{+/+}$ iPSC-cell derived neutrophils (day 28) is shown.

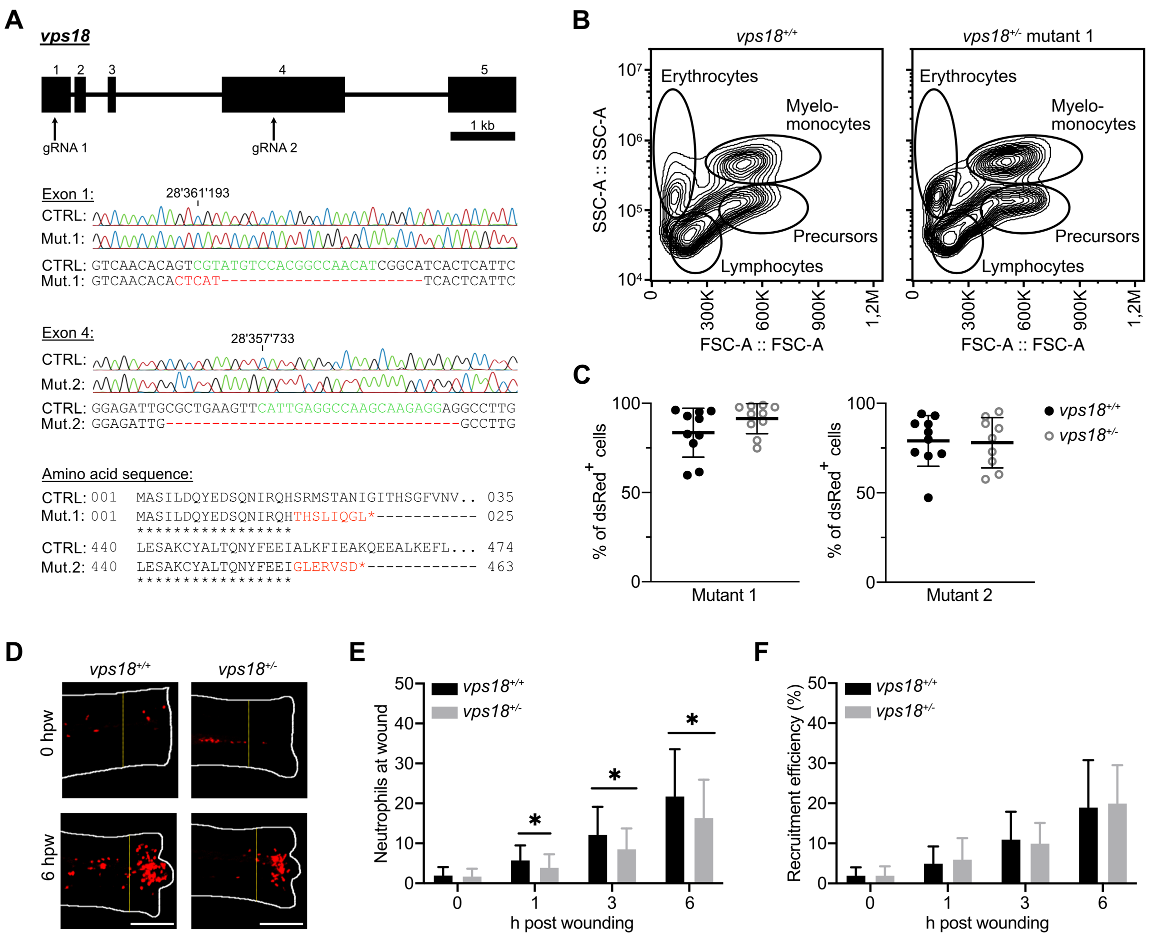


**Supplementary Fig. 7. *Vps18* mutant zebrafish larvae but not adult zebrafish exhibit reduced numbers of neutrophils.** (A) Upper panel: Schematic of zebrafish *vps18* gene and targeted exon 1 and 4 of gRNA 1 and 2, respectively. Middle panel: Sequencing traces and partial genomic sequence of *vps18*$^{+/+}$ (CTRL) and *vps18*$^{+/-}$ mutant (mut.) 1 and 2. Numbers indicate position within the gene. Target sequence of the gRNA, green. Nucleotides mutated or deleted in the mutants, red. Lower panel: Predicted amino acid sequence of mutants aligned to *vps18*$^{+/+}$ sequence. Identical (*) and altered (red) amino acids are indicated. (B) Scatter plot of single cell solution generated from whole kidney marrow (WKM) of adult *vps18*$^{+/+}$ and *vps18*$^{+/-}$ mutant 1 zebrafish. Gates indicate erythrocytes, lymphocytes, myelomonocytes and precursors of all lineages. (C) Quantification of dsRed$^{+}$ cells of myelomonocytes (100%) of WKM from (B) in adult *vps18*$^{+/+}$, *vps18*$^{+/-}$ mutant 1 and mutant 2 zebrafish. Mean ± SEM of ≥ 9 individual adult zebrafish of ≥ 2 independent experiments. (D) Representative maximum intensity projections of recruited neutrophils 0 and 6 h after tail fin transection of *vps18*$^{+/+}$ and *vps18*$^{+/-}$ mutant 1 zebrafish larvae at 5 dpf. Neutrophils, red. Area in which neutrophils were counted indicated by dashed lines. Scale bar, 200 µm. (E and F) Quantification of recruited neutrophils within the area depicted in (D) as absolute numbers (E) and recruitment efficiency (F).

Recruitment efficiency was calculated as ratio of recruited neutrophils/total neutrophil count. Mean ± SEM of ≥ 26 individual larvae of ≥ 3 independent experiments.
